# Cooperative NF-κB and Notch1 signaling promotes macrophage-mediated MenaINV expression in breast cancer

**DOI:** 10.1101/2023.01.03.522642

**Authors:** Camille L. Duran, George S. Karagiannis, Xiaoming Chen, Ved P. Sharma, David Entenberg, John S. Condeelis, Maja H. Oktay

## Abstract

Metastasis is a multistep process that leads to the formation of clinically detectable tumor foci at distant organs and frequently patient demise. Only a subpopulation of breast cancer cells within the primary tumor can disseminate systemically and cause metastasis. To disseminate, cancer cells must express MenaINV, an isoform of the actin-regulatory protein Mena encoded by the *ENAH* gene that endows tumor cells with transendothelial migration activity allowing them to enter and exit the blood circulation. We have previously demonstrated that MenaINV mRNA and protein expression is induced in cancer cells by macrophage contact. In this study, we discovered the precise mechanism by which macrophages induce MenaINV expression in tumor cells. We examined the promoter of the human and mouse *ENAH* gene and discovered a conserved NF-κB transcription factor binding site. Using live imaging of an NF-κB activity reporter and staining of fixed tissues from mouse and human breast cancer we further determined that for maximal induction of MenaINV in cancer cell NF-κB needs to cooperate with the Notch1 signaling pathway. Mechanistically, Notch1 signaling does not directly increase MenaINV expression, but it enhances and sustains NF-κB signaling through retention of p65, an NF-κB transcription factor, in the nucleus of tumor cells, leading to increased MenaINV expression. In mice, these signals are augmented following chemotherapy treatment and abrogated upon macrophage depletion. Targeting Notch1 signaling *in vivo* decreased NF-κB signaling and MenaINV expression in the primary tumor and decreased metastasis. Altogether, these data uncover mechanistic targets for blocking MenaINV induction that should be explored clinically to decrease cancer cell dissemination and improve survival of patients with metastatic disease.

## Introduction

Breast cancer is the second leading cause of cancer-related mortality in women in the US. Since the majority of breast cancer mortality is due to metastases, understanding the mechanisms that drive metastases is fundamental for the development of anti-metastatic therapies to improve the survival of patients with metastases.

The cell-biological program called “epithelial-to-mesenchymal transition” (EMT)(1, 2), during which cancer cells lose epithelial polarity and cell to cell cohesion(2, 3) is, in most instances, required for the onset of the metastatic cascade. The EMT program has been associated with heterotypic interactions of cancer cells with stromal and immune cells (e.g. macrophages), as well as with modified extracellular matrix, a hallmark of cancer progression and metastasis(2, 4, 5). During EMT, mRNAs encoding various proteins undergo alternative splicing(6), including mRNA for Mammalian enabled (Mena). EMT-induced alternative splicing of mRNA that encodes Mena, a protein involved in regulation of actin dynamics, results in a decrease of the non-metastatic isoform, Mena11a(6-9). However, the generation of dissemination-competent cancer cells requires an additional step: tumor cell-macrophage collisions, which lead to an increase in the expression of the MenaINV isoform(10). MenaINV enhances invasive cell motility(9, 11) and sensitizes cells to receptor tyrosine kinase (RTK) growth factors. These properties enable cancer cells to engage in a paracrine EGF-CSF1 signaling loop with tumor-associated macrophages (TAMs) and establish streaming migration with macrophages towards HGF-secreting endothelial cells (ECs)(12-15).

MenaINV expression in breast tumor cells is crucial for the formation of invadopodia, invasive protrusions required for cancer cell intravasation through portals on blood vessels called tumor microenvironment of metastasis (TMEM) doorways, and for extravasation at metastatic sites(16). Indeed, *in vivo* loss-of-function studies in *Mena* knockout mice, in which MenaINV expression is also eliminated, demonstrate a reduction in cell invasion, motility, intravasation, and metastatic dissemination in several mouse models(9, 17-19). Our recent studies demonstrate an increased density of MenaINV cancer cells, as well as cancer stem cells within 200 µm of TMEM doorways, where the most cancer cell-macrophage collisions occur, indicating that increased cancer cell-macrophage contact may be responsible for endowing cancer cells with both MenaINV and stem phenotypes.

We previously showed that MenaINV mRNA and protein expression in cancer cells involves macrophage-directed Notch1 signaling(10), and that macrophage expression of Jagged1 (a Notch signaling ligand) is critical for tumor cell intravasation(20). However, the promoter for the *ENAH* gene, which encodes Mena, does not contain binding sites for transcription factors in the Notch1 pathway. Thus, the mechanism of how macrophages induce expression of MenaINV, the protein required for tumor cell metastasis remains unidentified.

We have found, and independent reports confirm, that there is a κB binding site within the *ENAH* promoter, conserved from mouse to human(21). κB binding sites are used by transcription factors in the NF-κB signaling pathway, especially p65 (also known as RelA), to drive expression of target genes. There are numerous reports demonstrating Notch-mediated enhancement, and context-dependent activation, of NF-κB signaling in cancer(22-24). Thus, Notch1 signaling may activate the *ENAH* promoter indirectly via NF-κB signaling.

The NF-κB signaling pathway is known to play a major role in the progression of many cancers through promotion of processes such as EMT, proliferation, invasion, and resistance to cell death. NF-κB signaling can be activated by many factors produced within the TME, including proinflammatory cytokines such as TNFα and IL1β, growth factors, and oxidative stressors(25). It has become increasingly apparent that while NF-κB signaling can control a myriad of pro-invasive and pro-metastatic phenotypes, the downstream consequences of NF-κB activation are extraordinarily context dependent, and can, for example, enhance or inhibit apoptosis or tumor growth depending on the environment or stimulus(26-32).

As expression of MenaINV is essential for induction of an intravasation and extravasation-competent-competent phenotype in breast cancer cells which endows tumor cells the ability to metastasize, it is critical to determine if Notch1 signaling, caused by a juxtacrine, macrophage-tumor cell interaction, can promote NF-κB signaling, and subsequently contribute to increased MenaINV expression *in vivo*. Thus, here we investigated the hypothesis that macrophage-cancer cell interactions induce MenaINV expression in breast cancer cells through cooperation between Notch1 and NF-κB signaling. Understanding the mechanism by which tumor cells acquire MenaINV expression and its associated metastasis-inducing phenotypes is critical to aid the discovery of targetable signals to decrease metastatic burden and improve survival in breast cancer patients.

## Materials and Methods

### Cell lines and reagents

The MDA-MB-231 (231) human breast cancer cell line was purchased from ATCC, and the identity of the line was re-confirmed by STR profiling (Laragen Corp.), after expansion and passaging. The 6DT1 murine breast cancer cell line was generously provided by Dr. Lalage Wakefield, NCI. The MDA-MB-231 and 6DT1 cell lines were maintained in 10% FBS in DMEM with antibiotics. The BAC1.2F5 macrophage cell line was generously provided by Dr. Richard Stanley, Albert Einstein College of Medicine, and was maintained in 10% FBS in α-MEM with 3,000 units/ml CSF-1. All cells were maintained at 37°C in a 5% CO_2_ incubator, and were shown to be mycoplasma-free (Sigma LookOut Mycoplasma PCR detection kit, cat# MO0035-1KT). DAPT was reconstituted in 100% ethanol to a stock concentration of 20 mg/ml, aliquoted and stored at -20°C (Sigma, cat# D5942). DHMEQ (MedChemExpress, cat# HY-14645) was reconstituted in DMSO, aliquoted and stored at -20°C. C87 (Millipore Sigma, cat# 530796) was reconstituted in DMSO and stored at -80°C. SAHM1 (Millipore Sigma, cat# 491002) was reconstituted to 50 mg/ml in DMSO and stored at -20°C, clodronate liposomes (Encapsula Nano Sciences, cat# CLD-8901) were used as previously described(16). Jagged1 (Anaspec, cat# AS-61298) Jagged1 scrambled (Anaspec, cat# AS-64239) were reconstituted in DMSO, aliquoted and stored at -20°C and used at 80 µM/ml. Recombinant TNFα (Thermo Fisher, cat# PHC3015) was reconstituted at 0.1mg/ml in water, aliquoted and stored at -80°C, and used at 10 ng/ml. Active TGFβ was used at 5 and 10 ng/ml (abcam, cat# ab50036), LiCl was reconstituted in water, aliquoted and stored at -20°C, and used at 25 and 50 mM (Sigma Aldrich, cat# L9650), and Jagged1/2 blocking antibodies and IgG isotype control (Biolegend, cat#s 130902, 131001, 400902) were used at 20 µM/ml.

The MenaINV antibodies were generated by Covance, as previously described(10), and used at 0.25 µg/ml concentration for immunofluorescence staining. The p65 antibody used at 1:1000 for western blotting and staining (Cell Signaling Technology, cat# 8242S). Lamin A/C was used at 1:1000 for western blotting (Cell Signaling Technology, cat# 2032S), GAPDH was used at 1:10,000 for western blotting (Abcam, cat# ab8245), and Iba1 was used at 1:6,000 for staining (Wako, cat# 019-19741).

### Design of the NF-κB activity reporter

The GFP-p65 CDS was cloned out of the addgene plasmid (cat# 23255) by PCR, creating AfeI and PacI restriction enzyme cut sites, and ligated into the pT3-neo-Ef1a-GFP sleeping beauty vector from addgene (cat# 69134), cutting out the GFP sequence from the pT3-neo-Ef1a-GFP vector. Positive clones (pT3-neo-GFP-p65) were confirmed by sequencing. MDA-MB-231 and 6DT1 cells at 60% confluency were transiently transfected with 5.4 µg of pT3-neo-GFP-p65 and 0.6 µg of the transposase SB100 (addgene, cat# 34879) using 24 µl of Lipofectamine 2000 (Invitrogen). Stable MDA-MB-231/GFP-p65 and 6DT1/GFP-p65 cell lines were created by maintaining cells in 700 µg/ml G418, for 2 weeks. Expression of GFP-p65 was confirmed by western blotting and immunofluorescence staining using p65 and GFP antibodies and visual examination for GFP fluorescence. Cells were then flow sorted for top 90-95% of cells expressing GFP.

### Tumor cell and macrophage co-culture assay

MDA-MB-231 tumor cells (231) and BAC1.2F5 macrophages were co-cultured as previously described(10). In brief, 231 cells were seeded at 50% confluency in a 6-well plate and serum starved (0.5% FBS) overnight. The next morning, macrophages were seeded in the wells at a 1:5 ratio (231:macrophages), in media containing 0.5% FBS and 3000 units/ml CSF-1. At this point, any additional treatments or inhibitors were also added. Cells were allowed to incubate for 4 hours at 37°C in a 5% CO_2_ incubator before trypsinizing 231 tumor cells and making RNA or protein extracts.

### mRNA isolation and qPCR

Total RNA was isolated from tumor cells using RNA Mini Plus Kit (Qiagen, cat# 74134). cDNA was synthesized from 1µg total RNA using iScript cDNA synthesis (BioRad, cat# 1708891) following manufacturer’s instructions. Quantitative RT-PCR (qPCR) was performed with *Power* SYBR Green PCR Master Mix (applied biosystems, Thermo Fisher Scientific, cat# 4367659) using a QuantStudio 3 real-time PCR instrument (applied biosystems, Thermo Fisher Scientific). Expression of mRNA was normalized to human GAPDH expression levels as the endogenous control. The following primers were used: human GAPDH 5’- CGACCACTTTGTCAAGCTCA -3’, 5’- CCCTGTTGCTGTAGCCAAAT-3’; human MenaINV 5’- GATTCAAGACCATCAGGTTGTG - 3’, 5’- TACATCGCAAATTAGTGCTGTC -3’; human Hes1 5’- GTGAAGCACCTCCGGAAC -3’, 5’- GTCACCTCGTTCATGCACTC -3’; human IL-6 5’- AGCCACTCACCTCTTCAGAAC -3’, 5’- GCAAGTCTCCTCATTGAATCCAG -3’; mouse MenaINV 5’- AGAGGATGCCAATGTCTTCG -3’, 5’- TTAGTGCTGTCCTGCGTAGC -3’; and mouse GAPDH 5’- CATGTTCCAGTATGACTCCCTC -3’, 5’- GGCCTCACCCCATTTGATGT -3’.

### Live epifluorescence and analysis

GFP-p65 expressing tumor cells (MDA-MB-231/GFP-p65, 6DT1/GFP-p65) were seeded at 30% onto glass-bottom dishes (Mattek, cat# P35G-1.4-14-C) and serum starved overnight (0.5% FBS in DMEM). The next morning, the media was replaced with imaging media (0.5% FBS in L-15) and equilibrated at the heated (37°C) microscope for 2 hours. Cells were imaged live using an Olympus epifluorescence microscope with coolSNAP HQ2 CCD camera using a 40x objective. One 10x10 mosaic was captured and designated as time=0 and baseline GFP-p65 nuclear localization and the live imaging quickly paused. Any treatment (0.1-10 ng/ml TNFα, 80um Jagged1, or controls) were then added and imaging was immediately resumed and continued without interruption for 4 hours, taking an image of the same field approximately every 2.5 minutes. Timelapse movies were processed and analyzed using FIJI/ImageJ (NIH). To quantify the nuclear GFP-p65 localization over time, at time zero, a circular ROI was placed inside the nucleus and intensity of GFP signal was measured and designated as baseline GFP-p65 nuclear localization. The intensity of the GFP signal within the nuclear ROI was measured in each frame throughout the entire time course, moving the ROI (maintaining the same ROI size) only if the cell/nucleus moved in the frame. Forty-five cells were measured for each treatment, with three replicate dishes per treatment.

### Cell fractionation and western blotting

Cells were seeded into 6-well plates and serum starved overnight in 0.5% FBS in DMEM. Next morning, cells were treated with TNFα, Jagged1, or control (DMSO) at concentrations and times indicated in the figure legends. At the end of the treatment cells were trypsinized and cytoplasmic and nuclear fractions were separated and extracted using the NE-PER kit (Thermo-Fisher Scientific, cat# 78833) and stored at -80°C. Before western blotting, protein extracts were diluted at a 1:1 ratio in 2x laemmli sample buffer containing 2% 2-mercaptoethanol and boiled at

100°C for 5 minutes. Protein extracts were separated using a 10% sodium dodecyl sulfate polyacrylamide gel and transferred to immobilon polyvinylidene difluoride membranes (EMD Millipore). After blocking for one hour at room temperature in odyssey blocking buffer (LI-COR Biosciences), membranes were incubated with antibodies directed against p65 (1:1000, Cell Signaling Technologies, cat# 8242), Lamin A/C (1:1000, Cell Signaling Technologies, cat# 2032), or GAPDH (1:10,000, abcam, ab8245), rotating overnight at 4°C. Membranes were washed three times for five minutes with 0.1% Tween-20 in TBS before incubating for one hour with goat anti-mouse and goat anti-rabbit IRDye700CW-conjugated secondary antibodies (LI-COR Biosciences). Following three five-minute washes with 0.1% Tween-20 in TBS, membranes were scanned using a Classic Odyssey Infared Imager (LI-COR Biosciences). Quantitative analysis of images from three experiments was performed using FIJI/Image J software (NIH).

### Animal Models

All procedures were conducted in accordance with National Institutes of Health regulations and approved by the Albert Einstein College of Medicine Animal Use Committee. MDA-MB-231 cells were injected into the mammary fat pad of SCID mice (NCI) as previously described(33). Transgenic mice expressing the polyoma virus middle-T (PyMT) antigen under the control of the mammary tumor virus long terminal repeat (MMTV-LTR)(34) were bred in house and result in palpable tumors at approximately 6 weeks old. Patient derived xenograft (PDX) transplants of HT17 tumor chunks into SCID mice have been previously described(19, 35).

### In vivo treatments

#### Notch signaling inhibition in vivo using DAPT

Notch signaling inhibition *in vivo* using DAPT has been previously described(36). In brief, DAPT (Sigma-Aldrich, cat# D5942) was reconstituted in 100% ethanol to a stock concentration of 20 mg/ml, then further diluted in corn oil to a final concentration of 2 mg/ml. Eight-week-old PyMT mice bearing palpable tumors and separate cohort of SCID mice with tumors from orthotopically xenographed MDA-MB-231 cells were given daily intraperitoneal injections of 10 mg/kg DAPT or vehicle control (1:10 ethanol in corn oil) for 14 days. On day 15, the primary tumors were collected from the mice and fixed in 10% formalin. Mice were weighed on day 1 and day 15 to ensure no significant weight loss was suffered due to the DAPT treatment. Duodenums were stained using the Periodic acid-Schiff (PAS) staining and demonstrated an increase in goblet cell hyperplasia in the intestinal crypts (Supp. Fig. 6J(36)), consistent with successful Notch signaling inhibition *in vivo(37-39)*.

#### Macrophage depletion using clodronate liposomes

Macrophage depletion using clodronate liposomes *in vivo* has been previously described(16). Briefly, tumor bearing mice were treated with a 200 µl intraperitoneal injection of clodronate or PBS liposomes (Encapsula Nano Sciences, cat# CLD-8901) every other day for two weeks. After completion of the treatment, primary tumors were extracted from the mice and fixed in 10% formalin.

#### Paclitaxel and clodronate treatment

Paclitaxel treatment of mice *in vivo* has been previously described(19). Briefly, paclitaxel (Sigma-Aldrich) was reconstituted to a concentration of 10 mg/ml in 1:1 ethanol:cremophor-EL (Millipore, cat# 238470). Tumor bearing mice were treated *intravenously* with either 10 mg/kg paclitaxel (total of 200 µl) or 200 µl vehicle control (1:1 ethanol:cremophor-EL) every five days for two doses. Mice were randomly divided into four treatment groups: PBS liposomes and 1:1 ethanol:cremophor; PBS liposomes and paclitaxel; chlodronate liposomes and 1:1 ethanol:cremophor; and clodronate liposomes and paclitaxel. Treatment schemes are diagramed in Fig. 6A and Supp. Fig. 7A.

#### Tissue fixing, staining, and analysis

Following treatments described above, mice were sacrificed, and all mammary tumors were extracted and immersed in 10% formalin in a volume ratio of tumor to formalin of 1:7. Tissues were fixed for 24 to 48 hours and embedded in paraffin, then processed for histological examination. Paraffin blocks were cut into 10 µm thick sections and slides were deparaffinized by melting at 60°C in an oven equipped with a fan for 60 minutes, followed by 2x xylene treatment for 20 minutes. Slides were then rehydrated, and antigen retrieval was performed in 1 mM EDTA (pH 8.0) or 1x citrate buffer (pH 6.0) (Diagnostic BioSystems) at 97°C for 20 minutes in a conventional steamer. Endogenous peroxidase was blocked by using 0.3% hydrogen peroxide in water, followed by incubation of slides in a blocking buffer solution (10% FBS, 1% BSA, 0.0025% fish skin gelatin in 0.05% PBST) for 60 minutes at room temperature. Slides then were stained using the multiplex tyramide signal amplification (TSA) immunofluorescence assay, using the Perkin Elmer Opal 4-color Fluorescent IHC kit, according to the manufacturer’s instructions. The slides were stained with primary antibodies in sequence, against p65 (1:1000, Cell Signaling Technology, cat #8242S), Iba1 (1:6,000, Wako, cat# 019-19741), and MenaINV (1:1000, 0.25 μg/ml, see above). Slides were then washed three times in 0.05% PBST and incubated with secondary HRP-conjugated antibodies in appropriate sequence, including anti-rabbit and anti-chicken for 1 hour at room temperature. After washing three times with 0.05% PBST, slides were incubated with biotinylated tyramide (Perkin Elmer; Opal 4-color Fluorescent IHC kit) diluted at 1:50 in amplification buffer for 10 minutes. After washing, slides were incubated with spectral DAPI for 5 minutes and mounted with ProLong Gold antifade reagent (Life Technologies). The slides were imaged on the Pannoramic 250 Flash II digital whole slide scanner, using a 20x 0.75NA objective lens. Tissue suitable for scanning was automatically detected using intensity thresholding. Whole tissue images were uploaded in Pannoramic Viewer version 1.15.4 (3DHISTECH).

To measure p65 expression, p65 nuclear localization, MenaINV expression, and MenaINV expression associated with nuclear p65 in tissue section, a total of 10 different 40x fields were acquired per mouse, avoiding necrotic areas in the center of the tumor and the peritumoral stromal sheath at the rim of the tumor, which is devoid of tumor cells and infiltrated by inflammatory cells. The MenaINV, p65, and Iba1 channels were each thresholded just above background based upon intensity compared to the secondary antibody only control slide. Thresholding was achieved by only using linear methods, namely contrast adjustment.

#### Statistics

GraphPad Prism 7 and Excel were used to generate graphs/plots and for statistical hypothesis testing. Statistical significance was determined by either student’s *t*-test (normally distributed paired or unpaired dataset) or a one-way ANOVA with Tukey’s or Dunnett’s multiple comparisons test, as indicated in the figure legends. Statistical significance was defined as p-value < 0.05.

## Results

### NF-κB signaling mediates induction of MenaINV expression

We have previously found that MenaINV mRNA and protein expression are upregulated in breast cancer cells upon their direct cell contact with macrophages through Notch1 signaling(10). However, we now discovered that the promoter sequence for the *ENAH* gene does not contain RBP-J/CSL consensus binding sites, the transcription sites activated by Notch1 signaling. This finding indicates that Notch1 works in concert with other macrophage-mediated signals to induce MenaINV expression. We and others found consensus binding sites for transcription factors in NF-κB, Wnt, and TGFβ signaling pathways(21) (**Supp Fig. 1A**). Out of these three transcription binding sites only the κB site, located at -1070 and -850 in the *ENAH* promoter, is conserved across the species we examined: *H. sapiens, M. mulatta, M. musculus*, and *R. norvegicus* (**Supp Fig. 1A**). Neither TGFβ nor Wnt signaling induced MenaINV expression in human triple negative breast cancer cells MDA-MB-231 (231) in response to increasing doses of TGFβ and LiCl, activators of TGFβ and Wnt signaling, respectively(40, 41) (**Supp Fig. 1B &C**).

To test whether NF-κB signaling promotes MenaINV expression, we cultured 231 cells in the presence or absence of BAC1.2F5 macrophages with either TNFα, a potent activator of NF-κB signaling, or vehicle control. Co-culture of 231 cells with macrophages caused a 5-fold increase in MenaINV mRNA expression and treatment with TNFα led to a 1.7-fold increase in MenaINV expression. Under both conditions the increase in MenaINV mRNA was significant compared to the 231 cells cultured alone and treated with vehicle control (**Fig. 1A**). The addition of TNFα to the 231-macrophage co-culture did not enhance MenaINV expression beyond the level observed for 231-macrophage co-culture, indicating that the addition of TNFα is likely redundant to any signals provided by macrophages.

**Figure 1.**
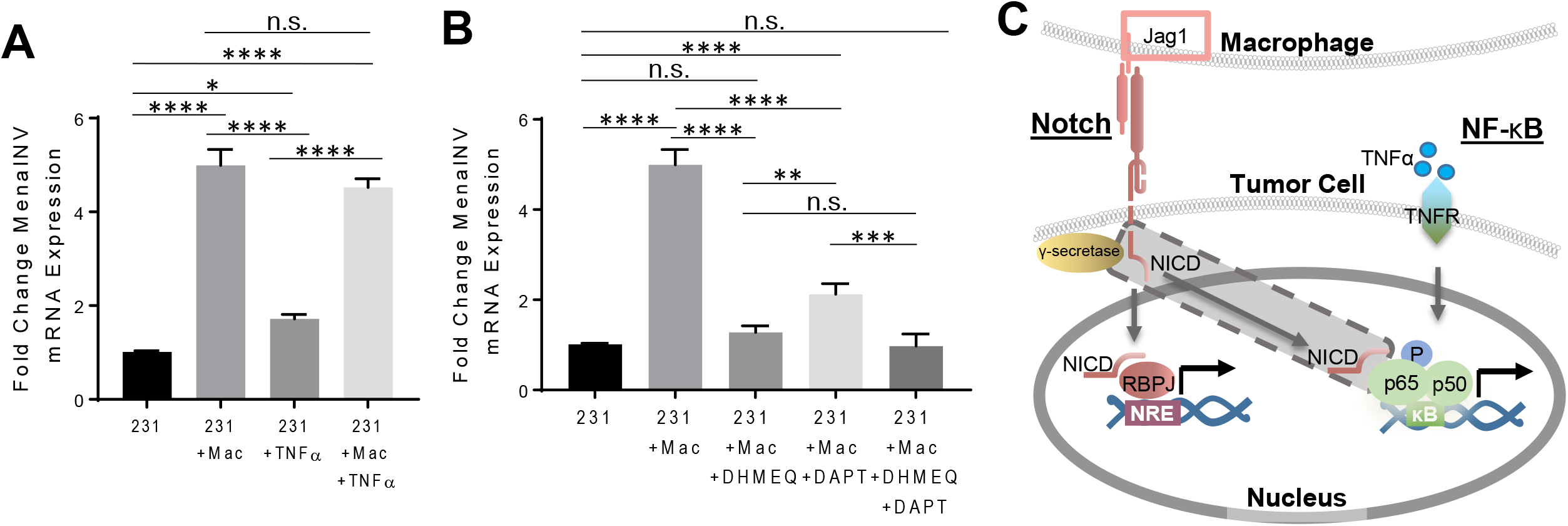
Macrophage-mediated induction of MenaINV expression via Notch and NF-κB cooperation. **(A)** MenaINV mRNA expression in MDA-MB-231 (231) cells co-cultured with or without BAC2.1F macrophages (Mac) and with or without 10ng/ml TNFα for 4 hours. **(B)** MenaINV mRNA expression in 231 cells co-cultured with or without Macs, NF-κB inhibitor (DHMEQ), or Notch/γ secretase inhibitor (DAPT) for 4 hours. **(C)** Model of potential Notch1 and NF-κB signaling “crosstalk” leading to enhanced transcriptional activity at the *ENAH* (Mena) promoter. The released Notch intracellular domain (NICD - shaded in gray) can bind to the transcription factors in the NF-κB signaling pathway and prevent their nuclear export, allowing for enhanced and sustained transcriptional activation of target genes and alternative splicing. The bars in (A) and (B) represent average fold change MenaINV mRNA compared to control (231 cells), +/-S.D. The data were analyzed using a one-way ANOVA with Tukey’s multiple comparisons test.*p<0.05, **p<0.01, ***p<0.001, ****p<0.0001, n.s.=not significant.

To test if NF-κB signaling is involved in the macrophage-induced MenaINV expression, we treated the 231-macrophage co-culture with the NF-κB signaling inhibitor, DHMEQ, and found the macrophage-induced increase in MenaINV mRNA expression in the 231 cells was abrogated back to the level observed for 231 cells cultured alone (**Fig. 1B**). We also found that the γ-secretase inhibitor, DAPT, which attenuates Notch signaling, only partially blocked the macrophage-induced increase in MenaINV mRNA expression. The addition of both DHMEQ and DAPT to the 231-macrophage co-culture brought MenaINV mRNA level back to that observed for 231 cells cultured alone (**Fig. 1B**). We ensured that the 231-macrophage co-culture was effectively activating Notch1 signaling by examining activation of Hes transcription, a transcriptional target of Notch1 signaling. Accordingly, we found that Hes mRNA levels were 3-fold higher in co-cultured cells compared to mono-cultured control 231 cells, and was abrogated when DAPT was added (**Supp Fig. 2**).

These results demonstrate that macrophage induced MenaINV expression in tumor cells requires the simultaneous activation of Notch1 and NF-κB signaling. Since NF-κB signaling is required but not sufficient to induce MenaINV expression to the levels achieved by the macrophage, we hypothesized that macrophages induce MenaINV expression in tumor cells through cooperation of Notch1 and NF-κB.

Cooperation between Notch1 and NF-κB leading to enhanced and prolonged signaling between the two pathways has previously been reported in other contexts(22). For example, upon Notch1 activation, the Notch intracellular domain (NICD) translocates to the nucleus where it can bind to the transcription factors of the NF-κB signaling pathway. This binding event blocks nuclear export of p65, causing nuclear retention of NF-κB transcription factors, and allowing for sustained and enhanced NF-κB signaling and transcription of NF-κB target genes(24). Thus, we hypothesized that macrophage-activated Notch1 enhances NF-κB signaling which increases MenaINV expression through prolonged nuclear retention of NF-κB transcription factor, p65 (**Fig. 1C**).

### Notch1 prolongs and sustains NF-κB signaling leading to MenaINV expression

To test the above hypothesis, we used an NF-κB reporter which allowed us to monitor, using live cell imaging, the activation of NF-κB signaling via direct visualization of p65 cellular localization. Briefly, we used a GFP sequence cloned upstream of the N-terminus of human p65 and cloned this GFP-p65 construct downstream of an EF-1α promoter in a sleeping beauty transposon vector. When NF-κB signaling is inactive, endogenous and GFP-p65 are retained in the cytosol, while upon NF-κB activation, endogenous and GFP-p65 are translocated to the nucleus (**Supp Fig. 3A**). We overexpressed the NF-κB reporter in 231 cells and 6DT1 mouse breast cancer carcinoma cells and tested several concentrations of TNFα in our system to activate NF-κB signaling. We determined that 10 ng/ml induces translocation of GFP-p65 into the nucleus within 30 minutes of the onset of treatment, while in the untreated cells, GFP-p65 was retained in the cytosol (**Supp Fig. 3B-D**).

To examine the levels of GFP-p65 compared to endogenous p65 and ensure there was no aberrant activation of NF-κB signaling in GFP-p65 overexpressing cells, we made nuclear and cytosolic extracts of 231 and 6DT1 GFP-p65 expressing cells treated with TNFα for 0, 10, and 30 minutes and then probed for p65 using western blotting. Cellular fractionation demonstrated the exogenous GFP-p65 was expressed at similar levels to endogenous p65 in both 231 and 6DT1 cells (**Supp Fig. 3E-H**). Although TNFα treated cells compared to untreated cells had significantly higher levels of nuclear p65, the amount of exogenous and endogenous p65, both nuclear and cytosolic, was similar in untreated and TNFα treated cells (**Supp Fig. 3E-H**). To ensure that NF-κB target genes were not aberrantly activated by the GFP-p65 reporter, we treated wild type and GFP-p65 expressing 231 cells with TNFα and measured induction of IL-6 mRNA expression, a cytokine potently expressed following NF-κB activation. We found that both wild type and GFP-p65 expressing 231 cells expressed similar levels of IL-6 mRNA following TNFα treatment, and the 231/GFP-p65 cells did not display any upregulation of IL-6 expression in untreated conditions, compared to wild type control 231 cells (**Supp Fig. 3I**). These results indicated that the NF-κB reporter was functional and could be used to monitor NF-κB signaling activation in both human and mouse mammary carcinoma cells.

To determine whether Notch1 signaling could potentiate NF-κB signaling, we incubated GFP-p65 expressing 231 and 6DT1 tumor cells with vehicle control, TNFα, Jagged1 (Notch1 ligand expressed on macrophages)(36), or both TNFα and Jagged1 combined for four hours and measured the intensity of green fluorescence signal in the nucleus over time. At time zero (t=0) we acquired one pre-treatment image (**Fig. 2A**), initiated one of the above treatments, and then continued time-lapse imaging for four hours (**Movies 1-4**). For the TNFα alone and TNFα+Jagged1 treatment groups, TNFα was added after the pre-treatment image and after 10 minutes of imaging, the TNFα-containing media was washed out and replaced with minimal media or Jagged1 containing media, respectively. Stills from the time lapse movies at 0, 17, and 240 minutes are shown in **Figure 2A** and the intensity of nuclear GFP-p65 signal at each time point is quantified in **Figure 2B**. In the vehicle control-treated cells, GFP-p65 was retained in the cytosol throughout the experiment, while in the TNFα treated cells, GFP-p65 robustly translocated into the nucleus within 17 minutes of the onset of treatment and two hours later shuttled back into the cytosol. In the Jagged1 treated cells, GFP-p65 shuttled into the nucleus very slowly over the course of four hours of imaging, never reaching the amplitude seen in the TNFα treated cells (**Fig. 2B**). The TNFα+Jagged1 treated cells demonstrated nuclear translocation of GFP-p65 at 17 minutes, as was seen in the TNFα-only treated cells, followed by nuclear retention of p65 throughout the four-hour time course (**Fig. 2A & B**). Similar results were obtained using 6DT1 GFP-p65 cells (**Supp Fig 4A, Movies 5-8**).

**Figure 2.**
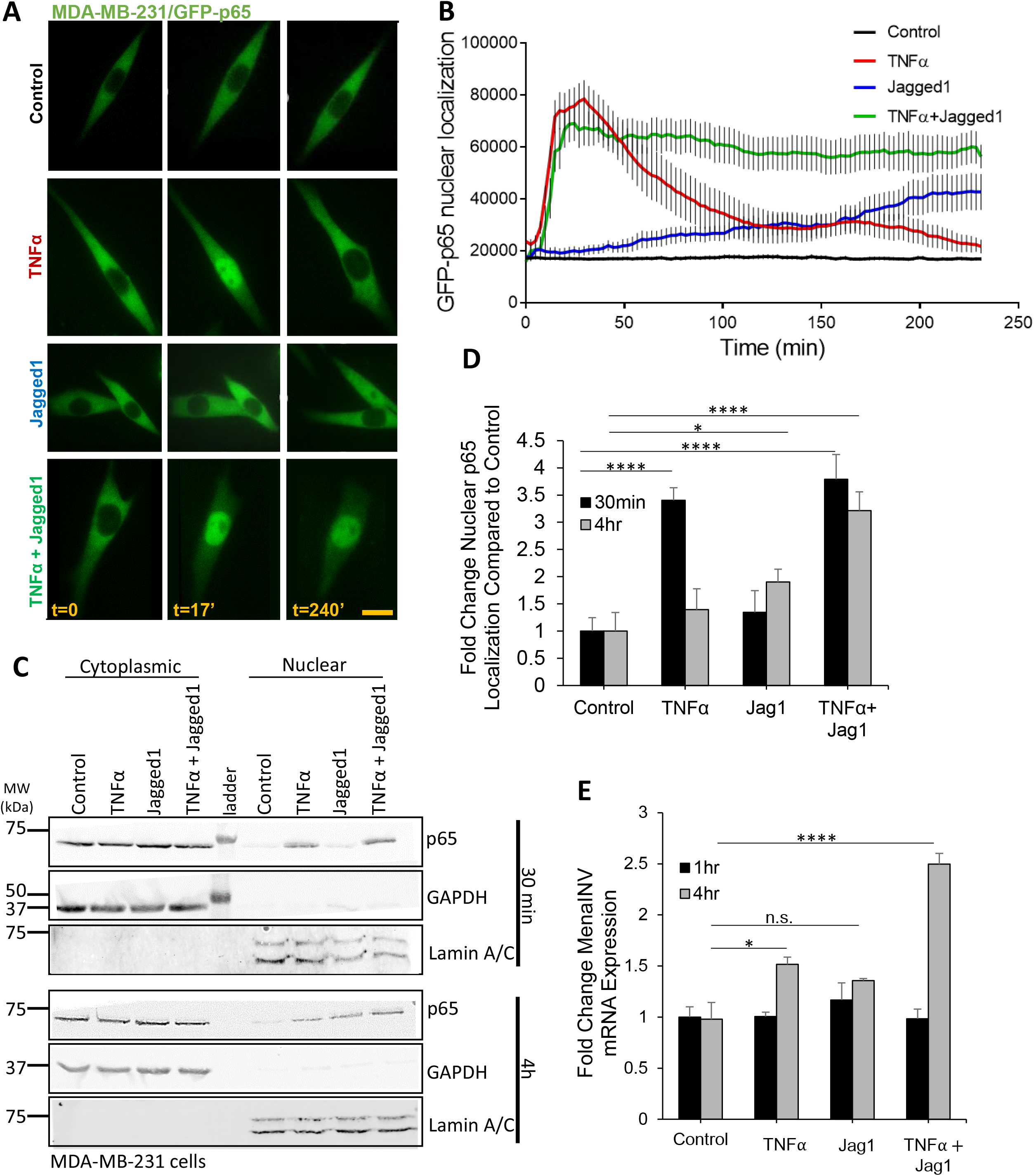
Notch1 enhances NF-κB signaling by sustaining p65 nuclear localization. **(A)** Stills from movies at 0, 17, and 240 minutes of MDA-MB-231/GFP-p65 cells treatment with vehicle, or 10 ng/ml human TNFα, or 80 µm Jagged1, or 10 ng/ml TNFα and 80 µm Jagged1. In all treatment groups with TNFα, the cells were treated for an initial 10 minutes, and then TNFα was washed out and replaced with minimal media, or with Jagged1 supplemented media. Cells were imaged live for 240 minutes using an EPI fluorescence microscope for the duration of the treatment, with one image captured every 2.5 minutes. Scale bar = 10 μm. **(B)** Quantification of normalized GFP-p65 nuclear localization over time from experiment in (A). **(C)** Western blot showing the amount of p65 in the cytoplasmic and nuclear fractions of wild type MDA-MB-231 cells treated for 30 minutes (upper blots) or 4 hours (lower blots) with vehicle, or 10 ng/ml TNFα, or 80 μm Jagged1, or 10 ng/ml TNFα and 80 μm Jagged1 (TNFα + Jagged1). In all treatment groups with TNFα, the cells were treated for an initial 10 minutes, and then TNFα was washed out and replaced with minimal media, or with Jagged1 supplemented media. (D) Quantification of western blots in (C) where the nuclear p65 signal was normalized to the lamin A/C signal. The graph shows the fold change in nuclear p65 signal for each treatment relative to the control treatment at both time points. **(E)** MenaINV mRNA expression in wild type MDA-MB-231 (231) cells treated as in (C) for 1 or 4 hours. Bars in (D) show average fold change MenaINV mRNA expression compared to Control at 1 or 4 hours. Data in (D) were analyzed using a one-way ANOVA with Tukey’s multiple comparisons test. *p<0.05, ****p<0.0001, n.s.=not significant.

To ensure that p65 nuclear translocation after treatment with TNFα and Jagged1 was not an artifact of the exogenously expressed GFP-p65, we treated wild type 231 cells with identical conditions and made nuclear and cytoplasmic extracts after 30 minutes and four hours of treatment and found a similar pattern of nuclear and cytosolic localization of endogenous p65 to that shown in the time lapse movies with GFP-p65 (**Fig. 2C & D**). To investigate if the above treatments lead to an increase in MenaINV expression, we treated wild type 231 cells accordingly and isolated mRNA after one and four hours of treatment. After one hour, none of the treatments had an effect on MenaINV mRNA expression. After four hours of treatment, TNFα alone caused a small but significant increase in MenaINV mRNA expression, Jagged1 alone had a slight but not significant increase in MenaINV mRNA expression, while TNFα+Jagged1 treatment led to a 2.5-fold increase in MenaINV mRNA expression (**Fig. 2E**). Similar results were obtained with 6DT1 cells (**Supp Fig 4B**). These data indicate that the treatment which resulted in the most robust and sustained activation of NF-κB signaling, as indicated by sustained nuclear p65 localization (**Fig. 2B-D**), also led to the most robust induction of MenaINV mRNA expression. In particular, the co-activation of Notch1 and NF-κB signalling (TNFα+Jagged1 treatment group) had a synergistic effect on MenaINV expression, compared to activation of each of the signaling pathway separately (**Fig. 2E**). Taken together these data indicate that the cooperation of Notch1 and NF-κB signaling is required for appreciable induction of MenaINV expression *in vitro*.

### Macrophage-mediated induction of MenaINV expression in tumor cells requires NF-κB and Notch1

To determine if the macrophage-mediated induction of MenaINV expression occurs specifically via TNFα and Notch1, we treated 231-macrophage co-cultures with more specific inhibitors, C87 and SAHM1, respectively. C87 is a small molecule inhibitor which directly binds to TNFα and blocks TNFα-induced NF-κB signaling(42). SAHM1 is a MAML1 inhibitor which prevents the NICD from binding to the transcriptional co-activator MAML1, leading to inhibition of Notch1 signaling downstream from receptor activation(43). While blocking TNFα activity with C87 almost completely abrogated the macrophage-induced expression of MenaINV, inhibition of MAML1 led to only partial reduction of MenaINV expression (**Fig. 3A**). Inhibition of both TNFα and MAML1 brought the macrophage-induced MenaINV expression to baseline levels observed when cancer cells were cultured without macrophages.

**Figure 3.**
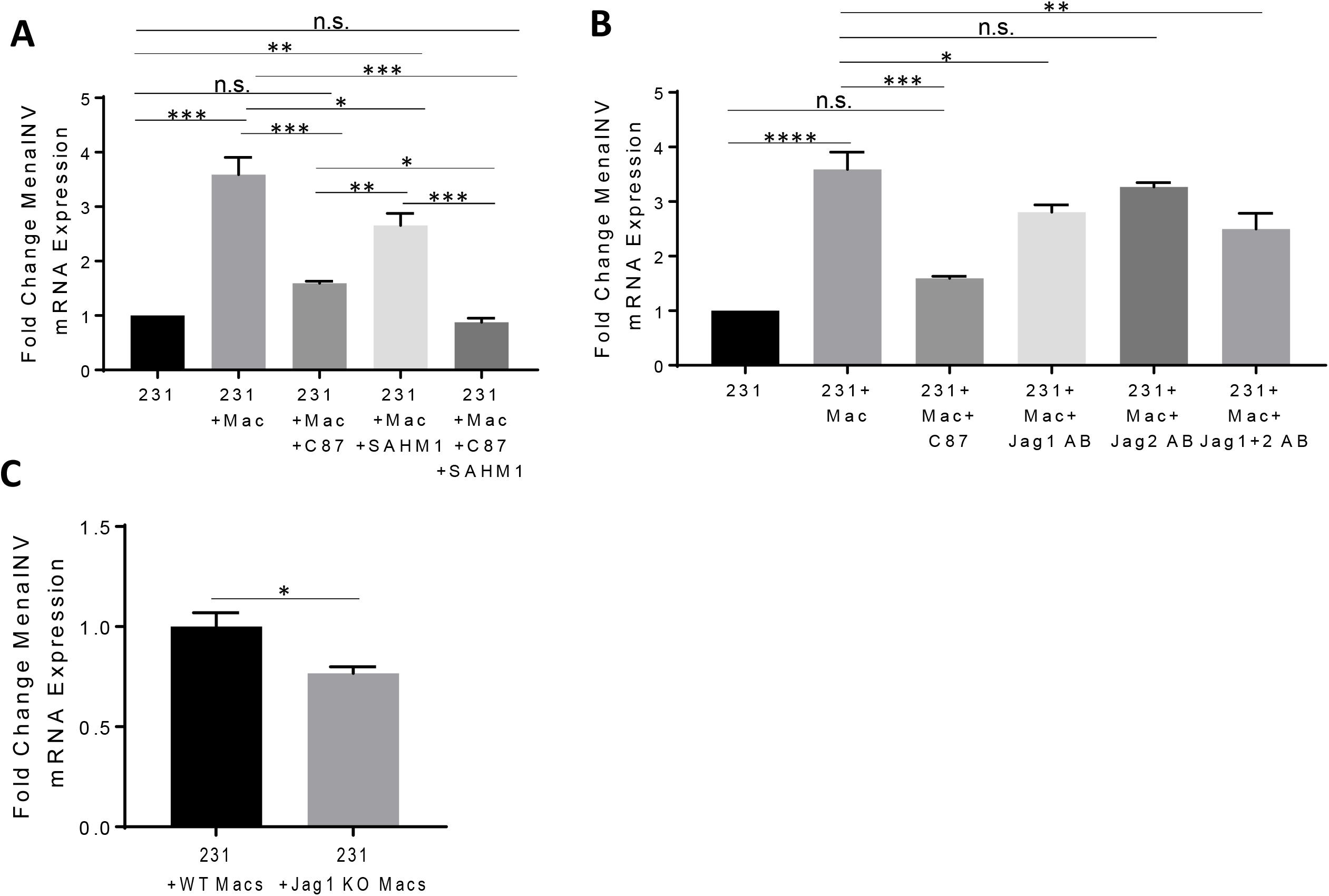
MenaINV expression in tumor cells induced by macrophages depends partially on TNFα mediated NF-κB signaling and Notch1 Jagged1 signaling. **(A)** MenaINV mRNA expression in MDA-MB-231 (231) cells co-cultured with or without BAC2.1F macrophages (Mac) and with or without C87 (TNFα inhibitor) or SAHM1 (MAML1 inhibitor) for 4 hours. **(B)** MenaINV mRNA expression in 231 cells co-cultured with or without Macs, TNFα inhibitor (C87), or Jag 1 or Jag2 blocking antibodies for 4 hours. Bars in (A) and (B) represent average fold change of MenaINV mRNA expression compared to control cells (231). **(C)** MenaINV mRNA expression in 231 cells co-cultured with wildtype (WT) or Jagged1 knockout BAC2.1F macrophages (Jag1 KO Macs). Bars in (A-C) represent average fold change of MenaINV mRNA expression compared to control cells (A and B: 231; C: 231 +WT Macs). Data were analyzed using a one-way ANOVA with Tukey’s multiple comparisons test. *p<0.05, **p<0.01, n.s.=not significant.

We next aimed to determine the specific Notch1 ligands on macrophages involved in the induction of MenaINV expression. While there are many Notch1 ligands, our recent study has found that the macrophages used in our co-culture experiments (BAC1.2F5), which show robust upregulation of MenaINV, primarily express Jagged1 and Jagged2, and an order of magnitude lower mRNA expression levels of Dll1, Dll2, and Dll4(36). Therefore, we focused our studies here on the role of Jagged1 and Jagged2 in the induction of MenaINV expression. We used Jagged1 and Jagged2 blocking antibodies to prevent Notch1 signaling activation in response to these macrophage-derived ligands. We found that blocking the Jagged1 ligand led to a modest but significant decrease in MenaINV expression compared to the tumor cell-macrophage co-cultured control group. Blocking the Jagged2 ligand did not significantly affect MenaINV mRNA expression. Blocking both Jagged1 and Jagged2 ligands together, did not lead to a further inhibition of MenaINV mRNA expression compared to blocking either ligand alone (**Fig. 3B**). This partial effect of blocking Jagged1/Notch1 signaling on MenaINV mRNA expression is consistent with the results seen with the more potent Notch1 inhibitors, DAPT and SAHM1. Taken together, these data indicate that macrophage-mediated induction of MenaINV expression occurs via TNFα and Jagged1. Furthermore, these data show that neither Notch1 nor NF-κB signaling on their own could fully account for the upregulation of MenaINV expression.

### Macrophage depletion decreases NF-κB signaling and MenaINV expression *in vivo*

We next wanted to determine whether macrophages are required for NF-κB mediated induction of MenaINV expression in cancer cells *in vivo*. We used two *in vivo* models of breast cancer previously generated in our laboratory: patient derived xenografts (PDX) from triple negative breast tumors (HT17) transplanted into *SCID* mice, and the autochthonous transgenic *MMTV-PyMT* transplantation model (PyMT), where a single spontaneously developed tumor is transplanted into the mammary fat pad of syngenic FVB mice(19, 35). The PyMT model fully recapitulates the entire breast cancer development and progression process(44). To deplete macrophages, we treated mice with clodronate liposomes (**Fig. 4A, Supp Fig. 5A**). Upon completion of treatment, we harvested the tumors and stained slides from the paraffin embedded tissues for the macrophage marker, Iba1 (to ensure our treatment decreased macrophage density in the primary tumor) (**Fig. 4B**), MenaINV, p65, and DAPI (**Fig. 4C and Supp Fig. 5B**). Treatment with clodronate liposomes, compared to control, decreased p65 expression in tumor cells (**Fig. 4D and Supp Fig. 5C**), and of the p65 that was expressed, less of it was localized in the nucleus of the tumor cells (**Fig. 4E and Supp Fig. 5D**). This indicates that macrophage depletion decreases expression as well as activation NF-κB signaling. Further, we found a corresponding decrease in MenaINV expression in tumors of clodronate-treated compared to control-treated, mice (**Fig. 4F and Supp Fig. 5E**). These data indicate that macrophage-mediated NF-κB activation is associated with MenaINV expression in tumor cells *in vivo*.

**Figure 4.**
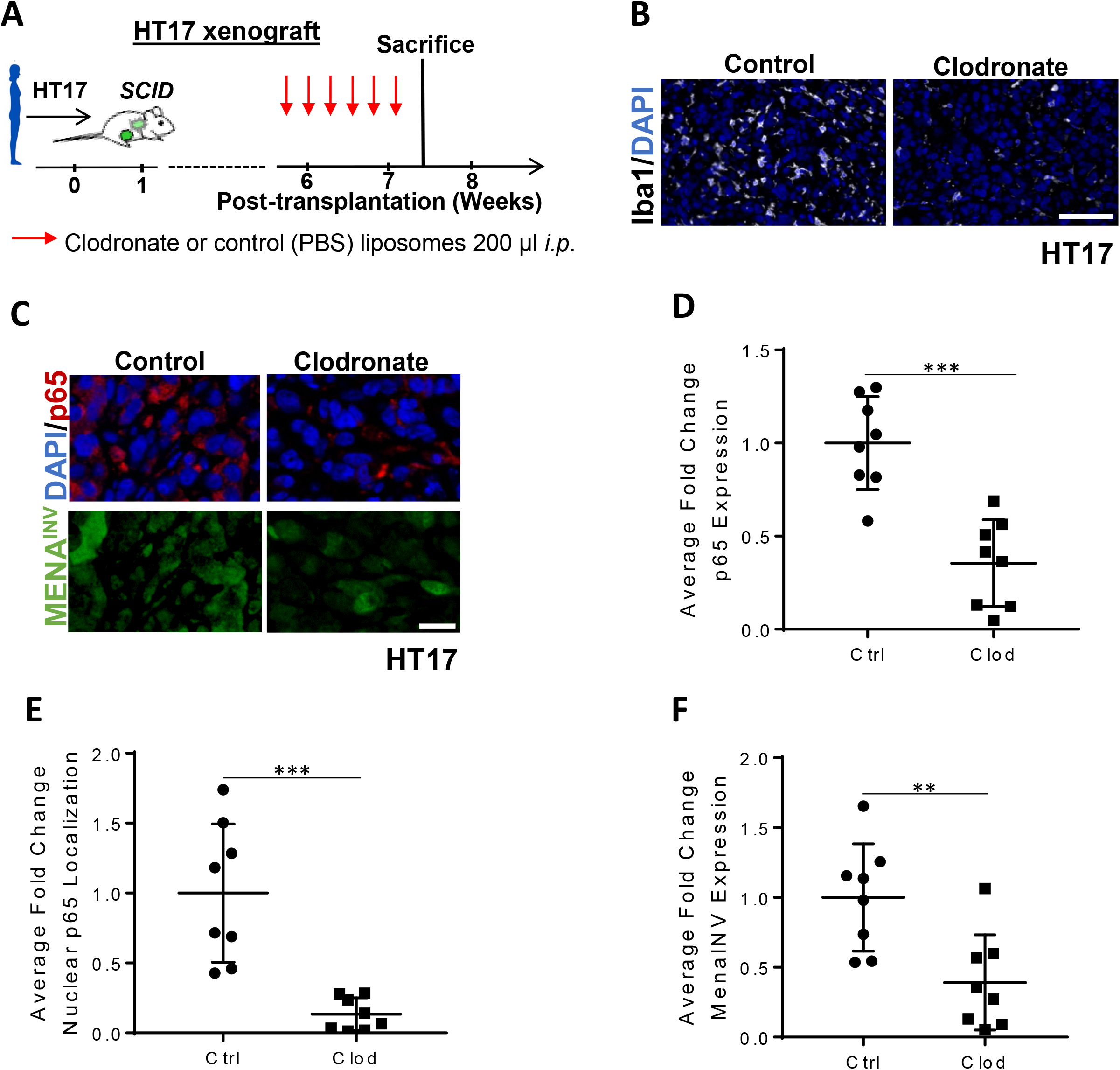
Macrophage depletion decreases NF-κB signaling and MenaINV expression in a PDX model in vivo. **(A)** Experimental design for macrophage depletion in patient derived xenografts (PDX) HT17 in SCID mice. *i*.*p*. = intraperitoneal. Red arrows indicate treatment days. **(B)** Immunofluorescence co-staining of HT17 xenografted in nude mice treated as outlined in (A) for the macrophage marker, Iba1 (white), **(C)** p65 (red), MenaINV (green) and nuclei (blue-DAPI). Blue and orange outlined sections demonstrate examples of what is quantified as primarily cytoplasmic (blue) or nuclear (orange) localization of p65. Scale bars = 100μm. **(D)** Quantification of average fold change in p65 expression from mice in (A). **(E)** Quantification of average fold change in p65 nuclear localization in PDX HT17 from mice treated as outlined in (A). Only p65 co-localized with the nuclear DAPI signal was quantified. **(F)** Quantification of average fold change MenaINV expression from PDX HT17 tumors treated as outlined in (A). Data in (D-F) were analyzed using a student’s *t*-test. **p<0.01.

### Inhibition of Notch signaling *in vivo* decreases activation of NF-κB and MenaINV expression in tumor cells

To determine whether inhibition of Notch1 signaling affects NF-κB activity and MenaINV expression *in vivo*, as observed *in vitro*, we treated mice bearing human breast cancer cell xenografts (MBA-MB-231 cells injected into the mammary fat pad) and PyMT(34) breast tumors with the γ-secretase inhibitor, DAPT, or control for two weeks (**Fig. 5A and Supp Fig. 6A**)(36). Upon completion of treatment, we harvested the tumors and stained them for p65, MenaINV, and DAPI (**Fig. 5B and Supp Fig. 6B**). We found a decrease in nuclear p65 (active NF-κB) in mice treated with DAPT compared to control mice in both models of breast cancer (**Fig. 5C and Supp Fig 6C**). Moreover, we observed a corresponding decrease in overall MenaINV expression in mice treated with DAPT, compared to control mice, in both models (**Fig. 5D and Supp Fig. 6D**). These results indicate that Notch1 inhibition *in vivo* decreases expression of MenaINV in an NF-κB dependent manner.

**Figure 5.**
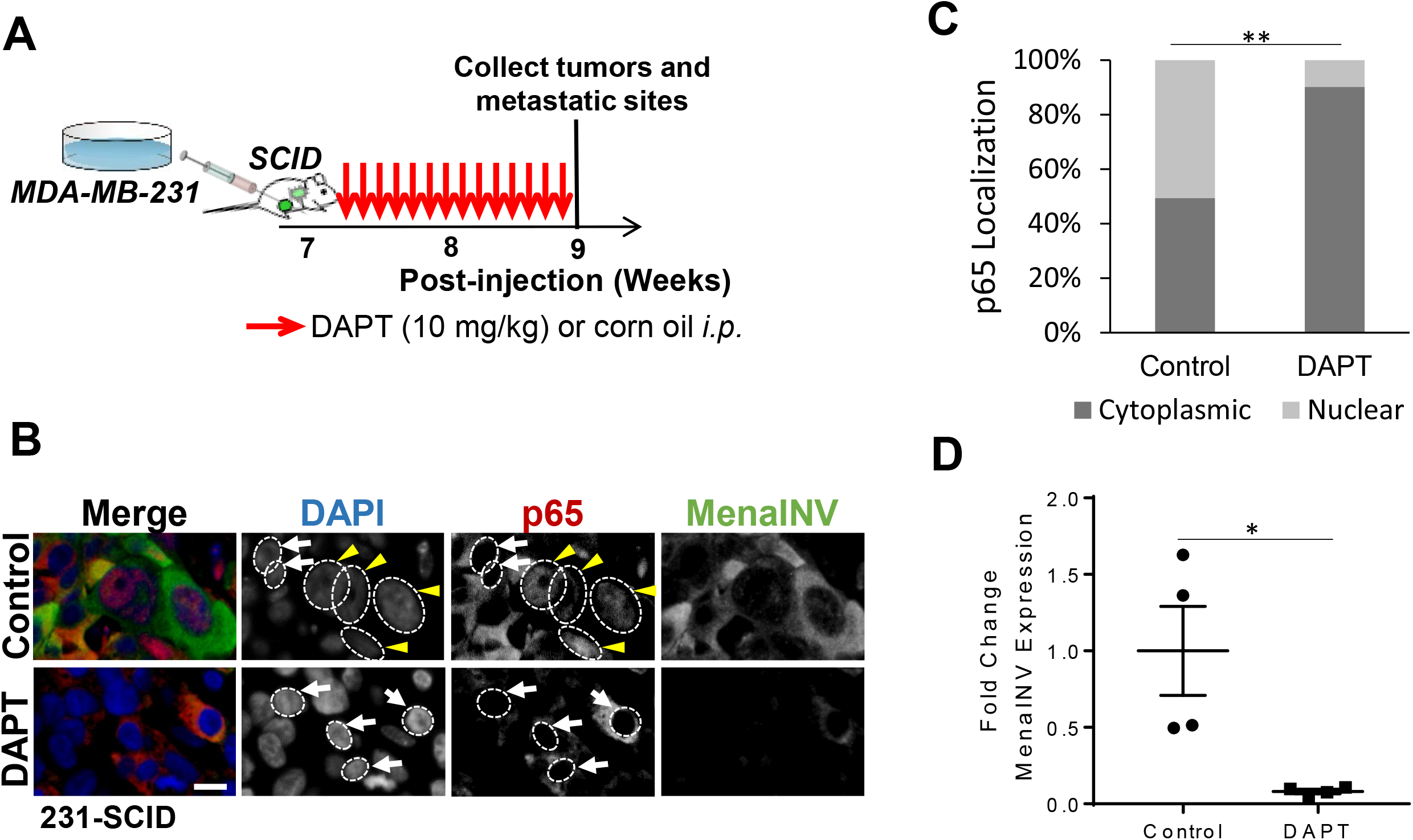
Inhibition of Notch1 signaling *in vivo* decreases activation of NF-κB signaling in MDA-MB-231 orthotropic injection model. **(A)** Schematic of DAPT treatment of *SCID* mice bearing orthotopically injected MDA-MB-231 tumor cells. Seven weeks post tumor cell injection mice were treated with 10 mg/kg DAPT or vehicle (corn oil) by *i*.*p*. every day for 14 days. Red arrows represent treatment days. **(B)** Immunofluorescence staining of primary tumor tissues sections for DAPI (nuclear stain, blue), p65 (red) and MenaINV (green). White dotted circles indicate nuclei in the DAPI and p65 channels. Yellow arrow heads denote nuclei with p65 positive stain (active NF-κB signaling), white arrowheads indicate nuclei without p65 positive staining (inactive NF-κB signaling). **(C)** Quantification of p65 localization (%cytoplasmic/nuclear) in tumor tissue from (B). **(D)** Quantification of average fold change in MenaINV expression compared to control mice from (B). Data in (C) and (D) were analyzed using a student’s *t*-test. *p<0.05, **p<0.01.

### Chemotherapy treatment enhances NF-κB activation and MenaINV expression through macrophage recruitment

We have previously shown that chemotherapy induces recruitment of macrophages into the tumor microenvironment, expression of MenaINV in transgenic and xenograft (human and mouse) mammary breast carcinoma models, and expression of MenaINV in residual breast cancer in patients after neoadjuvant treatment(19). We hypothesized that the chemotherapy-mediated increase in MenaINV expression occurs via macrophage recruitment and subsequent macrophage-mediated increase in NF-κB signaling. We tested this hypothesis by depleting the macrophages using clodronate in chemotherapy treated and untreated mice.

Briefly, mice bearing HT17 human PDXs or syngenic mouse PyMT tumors were treated with clodronate or control liposomes and either vehicle control (Ctrl) or paclitaxel (Ptx) as outlined in **Fig. 6A** and **Supp Fig. 7A**. Upon completion of treatment we compared the fold change in p65 nuclear localization (NF-κB activation) and MenaINV expression in tumors among the treatment groups (**Fig. 6B-D and Supp Fig. 7B-D**). Paclitaxel treatment significantly increased p65 nuclear localization (NF-κB activation) compared to vehicle control, while treatment with clodronate liposomes not only abrogated this increase, but also decreased nuclear p65 below the baseline of control animals which did not receive paclitaxel (**Fig. 6C and Supp Fig. 7C**). These findings indicate that macrophages are required for NF-κB activation in both chemotherapy treated and treatment-naïve animals. Furthermore, MenaINV expression in the tumor cells followed the same trend as the NF-κB signaling activation: paclitaxel, compared to control, increased MenaINV expression whereas clodronate abrogated paclitaxel-mediated induction of MenaINV expression as well as lowered MenaINV expression below the baseline observed in chemotherapy naïve animals (**Fig. 6D and Supp Fig 7D**).

**Figure 6.**
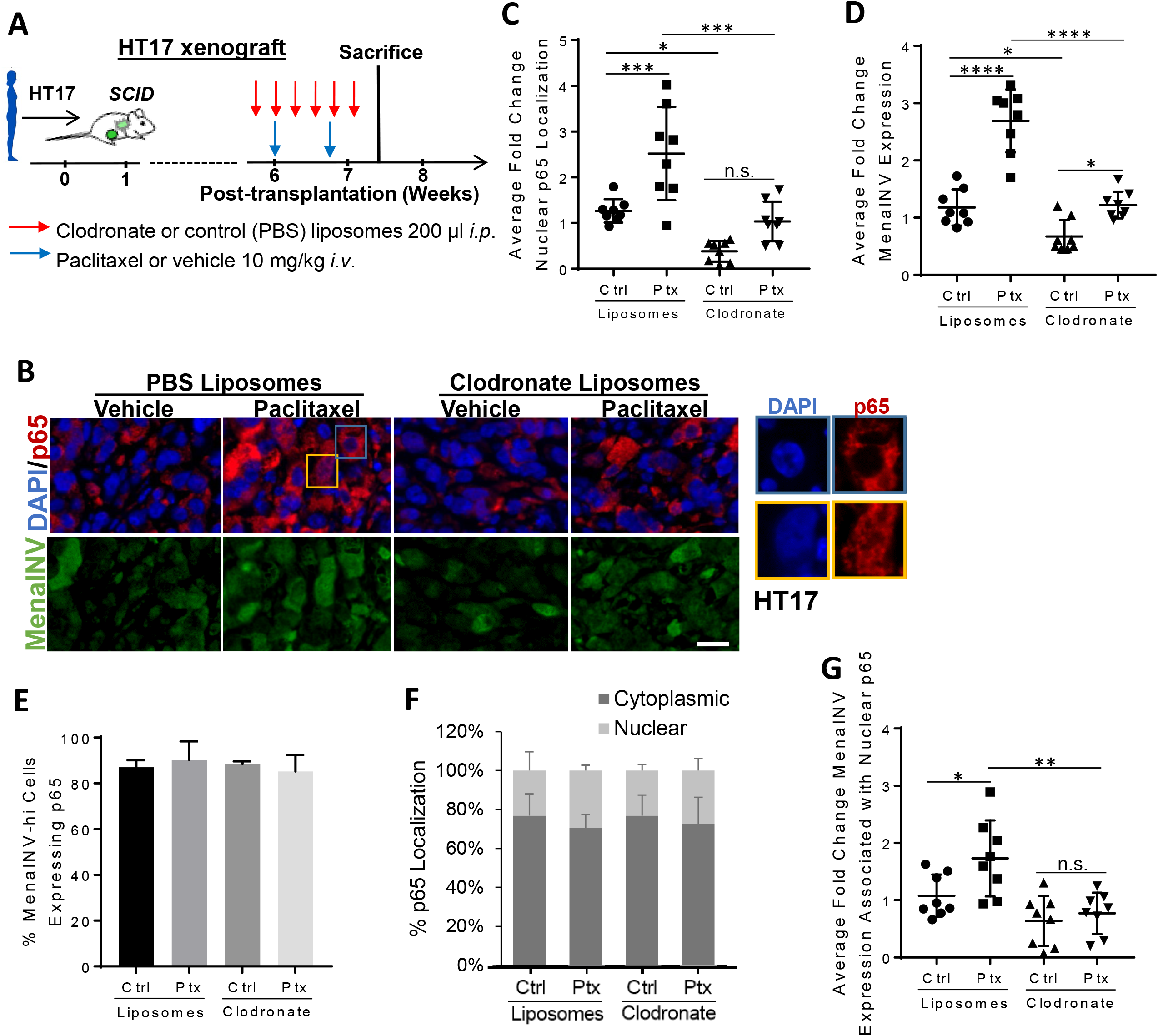
Chemotherapy treatment enhances NF-κB activation and MenaINV expression through macrophage recruitment in patient xenograft model. **(A)** Experimental design of chemotherapy and clodronate treatments in patient derived xenografts (PDX) HT17 in SCID mice. *i*.*p*. = intraperitoneal, *i*.*v*. = intravenous. **(B)** Immunofluorescence staining of primary breast tumor tissues from mice treated as outlined in (A) with DAPI (nuclear stain, blue), and antibodies recognizing p65 (red), and MenaINV (green). Blue and orange outlined sections are expanded below and demonstrate examples of what is quantified as primarily cytoplasmic (blue) or nuclear (orange) localization of p65 in HT17 tumor tissue. **(C)** Quantification of average fold change in p65 nuclear localization in treated primary tumors from (A) stained for p65 and DAPI. Only p65 which co-localized with the nuclear DAPI signal was quantified. **(D)** Quantification of average fold change in MenaINV expression in treated primary tumors from (A). **(E)** Quantification of the percentage of MenaINV-hi expressing tumor cells which also co-expressed p65 (regardless of cellular compartment localization), in primary tumor cells from treatments in (A). **(F)** Quantification of the localization (% cytoplasmic/nuclear) of p65 in MenaINV-hi expressing tumor cells from primary tumor cells treated in (A). **(G)** Quantification of average fold change MenaINV expression associated with nuclear p65 staining of primary tumors from (A) stained for MenaINV. Data in (C, D, and G) were analyzed using a one-way ANOVA with Tukey’s multiple comparisons test. *p<0.05, **p<0.01, ***p<0.001, n.s.=not significant.

To examine the relationship between p65 and MenaINV expressing cells, we measured whether tumor cells expressing MenaINV also express p65 in the same cell, and found that almost 90% of the MenaINV-hi expressing tumor cells also express p65, regardless of treatment (**Fig 6E**). Of the tumor cells which express both p65 and MenaINV-hi, we found that the majority (∼65-70%) of tumor cells express p65 in the cytoplasm, indicating that NF-κB signaling is not constitutively activated in these tumor cells under any treatment condition. (**Fig. 6F**).

To determine if active (nuclear p65) NF-κB signaling was associated with MenaINV expression, we measured the average fold change in MenaINV expression in cells where p65 was localized in the nucleus in treated mice compared to control mice (**Fig. 6G**). We found in the paclitaxel treatment group (where we had previously found the most robust NF-κB signalling activation), MenaINV was more highly expressed when p65 was nuclear, whereas in the clodronate treatment group (which has the lowest NF-κB activation), there was decreased MenaINV expression associated with nuclear p65. These results indicate that in the treatment group where NF-κB signaling is most robust and sustained, there is a concomitant upregulation of MenaINV expression in tumor cells. Similar results were obtained using the PyMT model of breast cancer (**Supp Fig 7E**). Taken together, these data demonstrate that macrophages are required for NF-κB activation and associated MenaINV expression *in vivo* in both chemotherapy-treated and chemotherapy naïve animals.

## Discussion

We discovered here that the specific mechanism by which tumor cells acquire swift and sustained expression of the metastasis-inducing protein, MenaINV, is via macrophage-mediated co-operative NF-κB and Notch1 signaling. Although previously found to be involved in the induction of MenaINV expression in response to macrophage and tumor cell contact, Notch1 signaling alone was unable to induce MenaINV expression (**Fig. 7A**). However, we determined that MenaINV can be induced by tumor-associated macrophages directly through macrophage-mediated activation of NF-κB, increasing expression of MenaINV by 1.5-fold (**Fig. 7B**). MenaINV expression can be further enhanced to 2.5-fold when Notch1 is activated in addition to NF-κB in tumor cells. Mechanistically, activation of Notch1 signaling in tumors cells by Jagged1-expressing macrophages leads to prolonged nuclear retention of the NF-κB transcription factor p65, and subsequent increase of MenaINV expression in tumor cells (**Fig. 7C)**. Importantly, we determined that the mechanism by which chemotherapy treatment enhances MenaINV expression occurs also through increased macrophage recruitment and subsequent cooperative Notch1 and NF-κB signaling in tumor cells.

**Figure 7.**
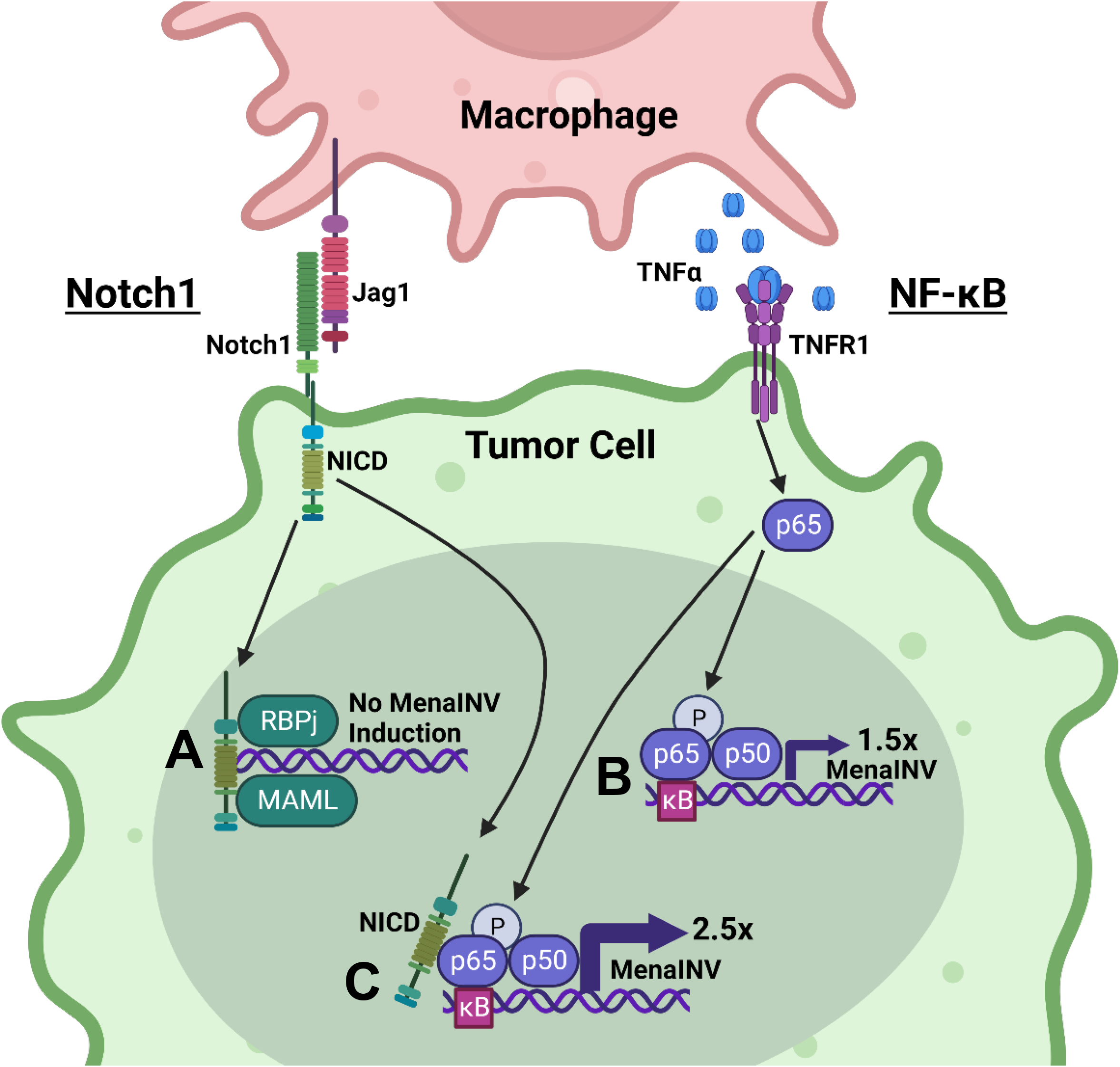
MenaINV expression in cancer cells is induced by macrophage-mediated co-operative NF-κB and Notch1 signaling. Juxtacrine and paracrine signaling between macrophages and tumor cells activate Notch1 and NF-κB pathways which co-operate to induce MenaINV expression in cancer cells. (**A**) Notch1 signaling alone does not induce MenaINV expression in tumor cells. (**B**) NF-κB signaling, activated by TNFα binding to the TNFR1 receptor, causes nuclear translocation of the transcription factor p65 and a 1.5-fold increase in MenaINV expression. (**C**) Notch1 and NF-κB signaling crosstalk to increase MenaINV expression further to 2.5-fold. Notch1 intracellular domain (NICD) enhances nuclear retention of NF-κB transcription factor p65 leading to sustained NF-κB signaling and induction of MenaINV expression. This mechanism of MenaINV induction is present *in vivo* and it explains previously observed increase in MenaINV expression upon in chemotherapy treatment(19). This detailed understanding of MenaINV induction in clinically relevant scenarios is needed for future development of combination therapies to improve survival of patients with breast cancer. Figure created with BioRender.com.

The precise mechanism of MenaINV induction in breast cancer cells shown here are of great translational significance for patients with metastases as previous studies have demonstrated that only cancer cells expressing the MenaINV isoform of the actin regulatory protein Mena, are capable of intravasating and metastasizing to secondary sites(12-15, 17, 19). MenaINV is required for formation of mature invadopodia which increase invasive and transendothelial migration capabilities of cancer cells(10, 18, 45). In addition, it was found that the expression of MenaINV occurs in macrophage-rich areas associated with TMEM doorways, increasing the likelihood that the MenaINV-expressing tumor cells will intravasate at TMEM doorways(36). Furthermore, MenaINV expressing tumor cells show a dramatically increased extravasation activity at distant sites, such as the lung, leading to highly efficient metastatic seeding(16).Therefore, discovering the mechanisms by which MenaINV expression is increased is important for understanding and targeting metastatic dissemination, which can occur not only from primary tumors, but also from metastatic foci resulting in overwhelming metastatic burden and patient demise(46-49).

Previous work demonstrated that macrophages induce MenaINV expression in tumor cells through Notch1 signaling(10). However, the Notch intracellular domain (NICD) which is cleaved from the intracellular portion of the receptor upon Notch1 activation, does not have DNA binding activity but acts as a transcriptional co-activator, along with MAML1 and RBP-J (CSL), to activate transcription of genes with RBP-J binding sites(50). We found that are no RBP-J binding sites within the *ENAH* promoter, and while we did not look at distant enhancer elements in this study, nonetheless, we determined that Notch1 could not induce MenaINV transcription directly. However, we and others, found κB sites within the *ENAH* promoter, conserved from mouse to human(21). Intriguingly, Notch1 and NF-κB signaling can crosstalk to enhance signaling of both signaling pathways(22-24). Our findings of macrophage-mediated direct induction of MenaINV expression by NF-κB, and indirect by Notch1 is supported by the fact that macrophages can provide stimuli to induce both Notch1 (Jagged1) and NF-κB (TNFα) signaling.

Shin et al. reported that NICD can bind to the NF-κB transcription factor, p65, which blocks p65 export from the nucleus, leading to sustained NF-κB signaling(24). Moreover, Field et al., found that an initial NF-κB signaling surge, followed by activation of a second NF-κB-independent signaling pathway, can lead to enhanced transcription of NF-κB target genes, and even increased levels of alternative transcripts(51). Consistent with these observations, we also found that Notch1 induces a prolonged nuclear retention of p65 and a subsequent surge in the expression of the MenaINV isoform of Mena, indicating that prolonged nuclear retention, through some still undiscovered mechanism, may affect alternative splicing. Intriguingly, there are also several p300 binding sites within the *ENAH* promoter which are binding sites for histone acetyltransferases. p65 has been found to promote strong activation of gene transcription following engagement with p300 and histone acetyltransferase activity(52).

Indeed, Notch1-mediated prolonged nuclear retention of p65 can explain our data showing that although NF-κB alone can induce MenaINV expression (1.5-fold), the level of induction is below the one achieved by macrophage-cancer cell contact (5-fold). The most robust activation of MenaINV expression was observed when Notch1 and NF-κB signaling were co-activated, by either macrophages or by specific Notch1 and NF-κB activating stimuli. Furthermore, NF-κB inhibitors blocked macrophage-mediated induction of MenaINV expression confirming that NF-κB signaling causes the initial increase in MenaINV expression, but that Notch1 signaling activation leads to the sustained NF-κB activity needed for the robust MenaINV expression.

Overall, these discoveries have important clinical implications because they indicate that increased intratumoral macrophage density, as encountered in certain clinical scenarios including high macrophage densities associated with TMEM doorways or inflammatory breast cancer, may affect disease progression. For example, several groups have shown that chemotherapy administration increases the density of tumor associated macrophages (TAMs)(19, 53, 54) and TMEM doorways(19). Thus, chemotherapy given pre-operatively to patients with more advanced disease may lead to an increase in intratumoral macrophage density and tumor cell dissemination via TMEM doorway activity(19). If chemotherapy fails to eradicate the tumor completely, an increased density of TAMs may subsequently enhance NF-κB signaling, which combined with Notch1 signaling, will increase MenaINV expression in residual tumor cells (**Fig. 7**). Our data explain the previously observed increase in MenaINV expression in the residual disease of some breast cancer patients who were treated with pre-operative (neoadjuvant) chemotherapy(19). Since chemotherapy is also given to patients with metastatic disease, one may speculate that macrophage recruitment and subsequent increase in MenaINV expression may occur in metastatic nodules as well. Moreover, it has been reported that chemotherapy not only increases MenaINV expression but also increases the density of TMEM doorways(19, 55) and potentially increases tumor cell dissemination via the blood circulation. Indeed, a recent study indicates that neoadjuvant chemotherapy in patients with early breast cancer leads to an increase of disseminated tumor cells in the bone marrow and subsequent worse overall survival(56). Therefore, the dismal five-year survival rate for breast cancer patients with metastatic disease of approximately 26% may be due to chemotherapy-induced cancer cell dissemination and increased cancer burden(54, 55, 57-60).

Paclitaxel is known to increase NF-κB signaling activation directly through binding to TLR4 receptors(61). We report here an additional mechanism of chemotherapy-mediated activation of NF-κB. This mechanism includes paclitaxel-mediated macrophage recruitment which leads to sustained NF-κB activation, potentially through Notch1. Given that MenaINV is critical for tumor cell invadopodium activation (which is involved in migration, invasion, and intravasation), the increase in NF-κB signaling activation and associated MenaINV expression following paclitaxel treatment observed here could explain previous studies that found that paclitaxel treatment increases circulating tumor cells (CTCs)(19).

The common therapies for advanced cancers, in addition to neoadjuvant chemotherapy, may include radiation. Interestingly, ionizing radiation is known to increase NF-κB signaling which may lead to NF-κB-mediated radiation resistance(62, 63). Thus, different treatment modalities for advanced cancer, while decreasing tumor mass, may inadvertently induce pro-metastatic changes in tumor microenvironment. These changes are characterized by macrophage recruitment, increased TMEM doorway density and activity(19), enhancement of NF-κB and Notch1 signaling in a subset of tumor cells leading to MenaINV-expression in cancer cells, followed by cancer cell dissemination through TMEM doorways, and ultimately increased metastatic burden.

The clinical use of Notch1 and NF-κB inhibitors were abandoned in the treatment of solid carcinoma due to toxicity when used systemically(64-66). Though the NF-κB signaling inhibitor bortezomib, a proteosome inhibitor, has been approved for treatment of multiple myeloma in patients who have failed two prior lines of therapy(67), there have not been many other successful uses of these inhibitors. As our knowledge and drug discovery platforms have improved over the last decades, it is important to revisit more specific inhibitors of Notch1 and NF-κB pathways as these signaling pathways may be enhanced by our current standard of care treatments.

## Conclusions

In summary, we have found that macrophages enhance expression of MenaINV, a pro-metastatic isoform of Mena, in breast cancer cells through Notch1-mediated prolongation of NF-κB activation. This macrophage-mediated sustained NF-κB signaling is seen *in vivo* and is enhanced by neoadjuvant chemotherapy. Thus, these findings underscore the need to further investigate combining inhibitors of Notch1 and NF-κB with chemotherapy to decrease chemotherapy-induced cancer cell dissemination and prolong survival of patients with advanced breast cancer.

## Declarations

### Ethics approval and consent to participate

All procedures were conducted in accordance with National Institutes of Health regulations and approved by the Albert Einstein College of Medicine Animal Use Committee.

## Consent for publication

Not applicable

## Availability of data and materials

Data sharing is not applicable to this article as no datasets were generated or analysed during the current study

## Competing interests

The authors declare that they have no competing interests

## Funding

This study was supported by grants from the NIH (R01 CA255153, F32 CA243350, K99 CA237851, an IRACDA fellowship, K12 GM102779), SIG OD019961, the Gruss-Lipper Biophotonics Center, the Integrated Imaging Program, The Evelyn Gruss-Lipper Charitable Foundation, and The Helen & Irving Spatz Foundation.

## Authors’ contributions

Conceptualization - CLD, MHO, JSC, DE

Methodology - CLD, GSK, XC

Formal Analysis - CLD

Software – DE

Investigation – CLD, GSK, XC, VPS

Writing - CLD, JSC, DE, MHO

Funding Acquisition – CLD, JSC, MHO, DE, GSK

Supervision - MHO, JSC, DE

All authors read and approved the final manuscript.

## Acknowledgements

We thank members of the Condeelis, Oktay, Entenberg, Cox, Segall, and Hodgson laboratories for helpful discussions. This study was supported by grants from the NIH (R01 CA255153, F32 CA243350, K99 CA237851, an IRACDA fellowship, K12 GM102779), SIG OD019961, the Gruss-Lipper Biophotonics Center, The Integrated Imaging Program, The Evelyn Gruss-Lipper Charitable Foundation, and The Helen & Irving Spatz Foundation.

## Notes

Conflicts of Interest: The authors declare no potential conflicts of interest.

### Competing Interest Statement

The authors have declared no competing interest.

## REFERENCES

1. Zenklusen D, Larson DR, Singer RH. Single-RNA counting reveals alternative modes of gene expression in yeast. Nat Struct Mol Biol. 2008;15(12):1263–71.

2. Kalluri R, Weinberg RA. The basics of epithelial-mesenchymal transition. J Clin Invest. 2009;119(6):1420–8.

3. Su S, Liu Q, Chen J, Chen J, Chen F, He C, et al. A positive feedback loop between mesenchymal-like cancer cells and macrophages is essential to breast cancer metastasis. Cancer Cell. 2014;25(5):605–20.

4. Karagiannis GS, Poutahidis T, Erdman SE, Kirsch R, Riddell RH, Diamandis EP. Cancer-associated fibroblasts drive the progression of metastasis through both paracrine and mechanical pressure on cancer tissue. Mol Cancer Res. 2012;10(11):1403–18.

5. Zhu Y, Knolhoff BL, Meyer MA, Nywening TM, West BL, Luo J, et al. CSF1/CSF1R blockade reprograms tumor-infiltrating macrophages and improves response to T-cell checkpoint immunotherapy in pancreatic cancer models. Cancer Res. 2014;74(18):5057–69.

6. Shapiro IM, Cheng AW, Flytzanis NC, Balsamo M, Condeelis JS, Oktay MH, et al. An EMT-driven alternative splicing program occurs in human breast cancer and modulates cellular phenotype. PLoS Genet. 2011;7(8):e1002218.

7. Gertler FB, Niebuhr K, Reinhard M, Wehland J, Soriano P. Mena, a relative of VASP and Drosophila Enabled, is implicated in the control of microfilament dynamics. Cell. 1996;87(2):227–39.

8. Pino MS, Balsamo M, Di Modugno F, Mottolese M, Alessio M, Melucci E, et al. Human Mena+11a isoform serves as a marker of epithelial phenotype and sensitivity to epidermal growth factor receptor inhibition in human pancreatic cancer cell lines. Clin Cancer Res. 2008;14(15):4943–50.

9. Roussos ET, Goswami S, Balsamo M, Wang Y, Stobezki R, Adler E, et al. Mena invasive (Mena(INV)) and Mena11a isoforms play distinct roles in breast cancer cell cohesion and association with TMEM. Clin Exp Metastasis. 2011;28(6):515–27.

10. Pignatelli J, Bravo-Cordero JJ, Roh-Johnson M, Gandhi SJ, Wang Y, Chen X, et al. Macrophage-dependent tumor cell transendothelial migration is mediated by Notch1/MenaINV-initiated invadopodium formation. Sci Rep. 2016;6:37874.

11. Eddy RJ, Weidmann MD, Sharma VP, Condeelis JS. Tumor Cell Invadopodia: Invasive Protrusions that Orchestrate Metastasis. Trends Cell Biol. 2017;27(8):595–607.

12. Patsialou A, Wyckoff J, Wang Y, Goswami S, Stanley ER, Condeelis JS. Invasion of human breast cancer cells in vivo requires both paracrine and autocrine loops involving the colony-stimulating factor-1 receptor. Cancer Res. 2009;69(24):9498–506.

13. Wang W, Mouneimne G, Sidani M, Wyckoff J, Chen X, Makris A, et al. The activity status of cofilin is directly related to invasion, intravasation, and metastasis of mammary tumors. J Cell Biol. 2006;173(3):395–404.

14. Wyckoff J, Wang W, Lin EY, Wang Y, Pixley F, Stanley ER, et al. A paracrine loop between tumor cells and macrophages is required for tumor cell migration in mammary tumors. Cancer Res. 2004;64(19):7022–9.

15. Leung E, Xue A, Wang Y, Rougerie P, Sharma VP, Eddy R, et al. Blood vessel endothelium-directed tumor cell streaming in breast tumors requires the HGF/C-Met signaling pathway. Oncogene. 2017;36(19):2680–92.

16. Borriello L, Coste A, Traub B, Sharma VP, Karagiannis GS, Lin Y, et al. Primary tumor associated macrophages activate programs of invasion and dormancy in disseminating tumor cells. Nat Commun. 2022;13(1):626.

17. Roussos ET, Wang Y, Wyckoff JB, Sellers RS, Wang W, Li J, et al. Mena deficiency delays tumor progression and decreases metastasis in polyoma middle-T transgenic mouse mammary tumors. Breast Cancer Res. 2010;12(6):R101.

18. Pignatelli J, Goswami S, Jones JG, Rohan TE, Pieri E, Chen X, et al. Invasive breast carcinoma cells from patients exhibit MenaINV- and macrophage-dependent transendothelial migration. Sci Signal. 2014;7(353):ra112.

19. Karagiannis GS, Pastoriza JM, Wang Y, Harney AS, Entenberg D, Pignatelli J, et al. Neoadjuvant chemotherapy induces breast cancer metastasis through a TMEM-mediated mechanism. Science Translational Medicine. 2017;9(397).

20. Cabrera RM, Mao SPH, Surve CR, Condeelis JS, Segall JE. A novel neuregulin - jagged1 paracrine loop in breast cancer transendothelial migration. Breast Cancer Res. 2018;20(1):24.

21. Urbanelli L, Massini C, Emiliani C, Orlacchio A, Bernardi G, Orlacchio A. Characterization of human Enah gene. Biochim Biophys Acta. 2006;1759(1-2):99–107.

22. Osipo C, Golde TE, Osborne BA, Miele LA. Off the beaten pathway: the complex cross talk between Notch and NF-kappaB. Lab Invest. 2008;88(1):11–7.

23. Guan E, Wang J, Laborda J, Norcross M, Baeuerle PA, Hoffman T. T cell leukemia-associated human Notch/translocation-associated Notch homologue has I kappa B-like activity and physically interacts with nuclear factor-kappa B proteins in T cells. J Exp Med. 1996;183(5):2025–32.

24. Shin HM, Minter LM, Cho OH, Gottipati S, Fauq AH, Golde TE, et al. Notch1 augments NF-kappaB activity by facilitating its nuclear retention. EMBO J. 2006;25(1):129–38.

25. DiDonato JA, Mercurio F, Karin M. NF-kappaB and the link between inflammation and cancer. Immunol Rev. 2012;246(1):379–400.

26. Greten FR, Eckmann L, Greten TF, Park JM, Li ZW, Egan LJ, et al. IKKbeta links inflammation and tumorigenesis in a mouse model of colitis-associated cancer. Cell. 2004;118(3):285–96.

27. Maeda S, Kamata H, Luo JL, Leffert H, Karin M. IKKbeta couples hepatocyte death to cytokine-driven compensatory proliferation that promotes chemical hepatocarcinogenesis. Cell. 2005;121(7):977–90.

28. Dajee M, Lazarov M, Zhang JY, Cai T, Green CL, Russell AJ, et al. NF-kappaB blockade and oncogenic Ras trigger invasive human epidermal neoplasia. Nature. 2003;421(6923):639–43.

29. van Hogerlinden M, Rozell BL, Ahrlund-Richter L, Toftgard R. Squamous cell carcinomas and increased apoptosis in skin with inhibited Rel/nuclear factor-kappaB signaling. Cancer Res. 1999;59(14):3299–303.

30. Nelson DE, Ihekwaba AE, Elliott M, Johnson JR, Gibney CA, Foreman BE, et al. Oscillations in NF-kappaB signaling control the dynamics of gene expression. Science. 2004;306(5696):704–8.

31. Ashall L, Horton CA, Nelson DE, Paszek P, Harper CV, Sillitoe K, et al. Pulsatile stimulation determines timing and specificity of NF-kappaB-dependent transcription. Science. 2009;324(5924):242–6.

32. Sung MH, Salvatore L, De Lorenzi R, Indrawan A, Pasparakis M, Hager GL, et al. Sustained oscillations of NF-kappaB produce distinct genome scanning and gene expression profiles. PLoS One. 2009;4(9):e7163.

33. Patsialou A, Bravo-Cordero JJ, Wang Y, Entenberg D, Liu H, Clarke M, et al. Intravital multiphoton imaging reveals multicellular streaming as a crucial component of in vivo cell migration in human breast tumors. Intravital. 2013;2(2):e25294.

34. Guy CT CR, Muller WJ. Induction of mammary tumors by expression of polyomavirus middle T oncogene: a transgenic mouse model for metastatic disease. Molecular and cellular biology. 1992;12(3).

35. Patsialou A, Wang Y, Lin J, Whitney K, Goswami S, Kenny PA, et al. Selective gene-expression profiling of migratory tumor cells in vivo predicts clinical outcome in breast cancer patients. Breast Cancer Res. 2012;14(5):R139.

36. Sharma VP, Tang B, Wang Y, Duran CL, Karagiannis GS, Xue EA, et al. Live tumor imaging shows macrophage induction and TMEM-mediated enrichment of cancer stem cells during metastatic dissemination. Nat Commun. 2021;12(1):7300.

37. Shah MM, Zerlin M, Li BY, Herzog TJ, Kitajewski JK, Wright JD. The role of Notch and gamma-secretase inhibition in an ovarian cancer model. Anticancer Res. 2013;33(3):801–8.

38. Milano J, McKay J, Dagenais C, Foster-Brown L, Pognan F, Gadient R, et al. Modulation of notch processing by gamma-secretase inhibitors causes intestinal goblet cell metaplasia and induction of genes known to specify gut secretory lineage differentiation. Toxicol Sci. 2004;82(1):341–58.

39. Sjolund J, Johansson M, Manna S, Norin C, Pietras A, Beckman S, et al. Suppression of renal cell carcinoma growth by inhibition of Notch signaling in vitro and in vivo. J Clin Invest. 2008;118(1):217–28.

40. Zhang X, Chen Q, Song H, Jiang W, Xie S, Huang J, et al. MicroRNA-375 prevents TGF-β-dependent transdifferentiation of lung fibroblasts via the MAP2K6/P38 pathway. Mol Med Rep. 2020;22(3):1803–10.

41. Clément-Lacroix P, Ai M, Morvan F, Roman-Roman S, Vayssière B, Belleville C, et al. Lrp5-independent activation of Wnt signaling by lithium chloride increases bone formation and bone mass in mice. Proc Natl Acad Sci U S A. 2005;102(48):17406–11.

42. Ma L, Gong H, Zhu H, Ji Q, Su P, Liu P, et al. A novel small-molecule tumor necrosis factor alpha inhibitor attenuates inflammation in a hepatitis mouse model. J Biol Chem. 2014;289(18):12457–66.

43. Moellering RE, Cornejo M, Davis TN, Del Bianco C, Aster JC, Blacklow SC, et al. Direct inhibition of the NOTCH transcription factor complex. Nature. 2009;462(7270):182–8.

44. Lin EY, Jones JG, Li P, Zhu L, Whitney KD, Muller WJ, et al. Progression to malignancy in the polyoma middle T oncoprotein mouse breast cancer model provides a reliable model for human diseases. Am J Pathol. 2003;163(5):2113–26.

45. Roussos ET, Balsamo M, Alford SK, Wyckoff JB, Gligorijevic B, Wang Y, et al. Mena invasive (MenaINV) promotes multicellular streaming motility and transendothelial migration in a mouse model of breast cancer. J Cell Sci. 2011;124(Pt 13):2120–31.

46. Borriello L, Condeelis J, Entenberg D, Oktay MH. Breast Cancer Cell Re-Dissemination from Lung Metastases-A Mechanism for Enhancing Metastatic Burden. J Clin Med. 2021;10(11).

47. Nguyen DX, Bos PD, Massague J. Metastasis: from dissemination to organ-specific colonization. Nat Rev Cancer. 2009;9(4):274–84.

48. Kim MY, Oskarsson T, Acharyya S, Nguyen DX, Zhang XH, Norton L, et al. Tumor self-seeding by circulating cancer cells. Cell. 2009;139(7):1315–26.

49. Zhang W, Bado IL, Hu J, Wan YW, Wu L, Wang H, et al. The bone microenvironment invigorates metastatic seeds for further dissemination. Cell. 2021;184(9):2471-86.e20.

50. Borggrefe T, Oswald F. The Notch signaling pathway: transcriptional regulation at Notch target genes. Cell Mol Life Sci. 2009;66(10):1631–46.

51. Field JT, Martens MD, Mughal W, Hai Y, Chapman D, Hatch GM, et al. Misoprostol regulates Bnip3 repression and alternative splicing to control cellular calcium homeostasis during hypoxic stress. Cell Death Discov. 2018;4:37.

52. Vanden Berghe W, De Bosscher K, Boone E, Plaisance S, Haegeman G. The nuclear factor-kappaB engages CBP/p300 and histone acetyltransferase activity for transcriptional activation of the interleukin-6 gene promoter. J Biol Chem. 1999;274(45):32091–8.

53. Hughes R, Qian BZ, Rowan C, Muthana M, Keklikoglou I, Olson OC, et al. Perivascular M2 Macrophages Stimulate Tumor Relapse after Chemotherapy. Cancer Res. 2015;75(17):3479–91.

54. DeNardo DG, Brennan DJ, Rexhepaj E, Ruffell B, Shiao SL, Madden SF, et al. Leukocyte complexity predicts breast cancer survival and functionally regulates response to chemotherapy. Cancer Discov. 2011;1(1):54–67.

55. Chang YS, Jalgaonkar SP, Middleton JD, Hai T. Stress-inducible gene Atf3 in the noncancer host cells contributes to chemotherapy-exacerbated breast cancer metastasis. Proc Natl Acad Sci U S A. 2017;114(34):E7159–E68.

56. Volmer L, Koch A, Matovina S, Dannehl D, Weiss M, Welker G, et al. Neoadjuvant Chemotherapy of Patients with Early Breast Cancer Is Associated with Increased Detection of Disseminated Tumor Cells in the Bone Marrow. Cancers (Basel). 2022;14(3).

57. Shaked Y. The pro-tumorigenic host response to cancer therapies. Nat Rev Cancer. 2019.

58. Karagiannis GS, Condeelis JS, Oktay MH. Chemotherapy-Induced Metastasis: Molecular Mechanisms, Clinical Manifestations, Therapeutic Interventions. Cancer Res. 2019;79(18):4567–76.

59. Daenen LG, Houthuijzen JM, Cirkel GA, Roodhart JM, Shaked Y, Voest EE. Treatment-induced host-mediated mechanisms reducing the efficacy of antitumor therapies. Oncogene. 2014;33(11):1341–7.

60. Shaked Y, Henke E, Roodhart JM, Mancuso P, Langenberg MH, Colleoni M, et al. Rapid chemotherapy-induced acute endothelial progenitor cell mobilization: implications for antiangiogenic drugs as chemosensitizing agents. Cancer Cell. 2008;14(3):263–73.

61. Ran S. The Role of TLR4 in Chemotherapy-Driven Metastasis. Cancer Res. 2015;75(12):2405–10.

62. Lee SJ, Dimtchev A, Lavin MF, Dritschilo A, Jung M. A novel ionizing radiation-induced signaling pathway that activates the transcription factor NF-kappaB. Oncogene. 1998;17(14):1821–6.

63. Li X, Chen F, Zhu Q, Ding B, Zhong Q, Huang K, et al. Gli-1/PI3K/AKT/NF-kB pathway mediates resistance to radiation and is a target for reversion of responses in refractory acute myeloid leukemia cells. Oncotarget. 2016;7(22):33004–15.

64. Gilmore TD, Herscovitch M. Inhibitors of NF-kappaB signaling: 785 and counting. Oncogene. 2006;25(51):6887–99.

65. Ryeom SW. The cautionary tale of side effects of chronic Notch1 inhibition. J Clin Invest. 2011;121(2):508–9.

66. Begalli F, Bennett J, Capece D, Verzella D, D’Andrea D, Tornatore L, et al. Unlocking the NF-κB Conundrum: Embracing Complexity to Achieve Specificity. Biomedicines. 2017;5(3).

67. Paramore A, Frantz S. Bortezomib. Nature Reviews Drug Discovery. 2003;2(8):611–2.

