## Supplemental Figures and Legends for "Cooperative NF-κB and Notch1 signaling promotes macrophage-mediated MenaINV expression in breast cancer"

**Supplemental Figure 1.**

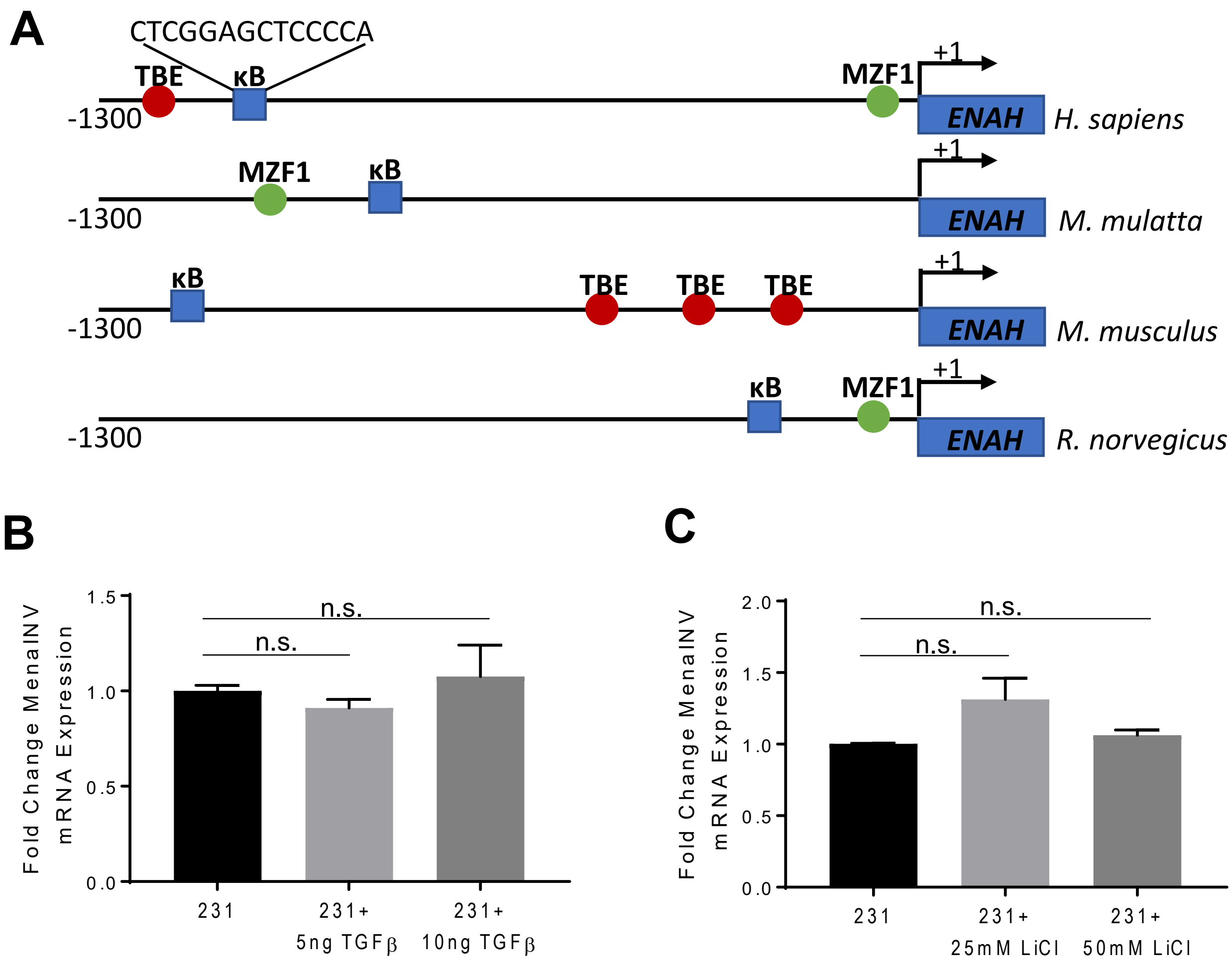

**Supplemental Figure 2.**

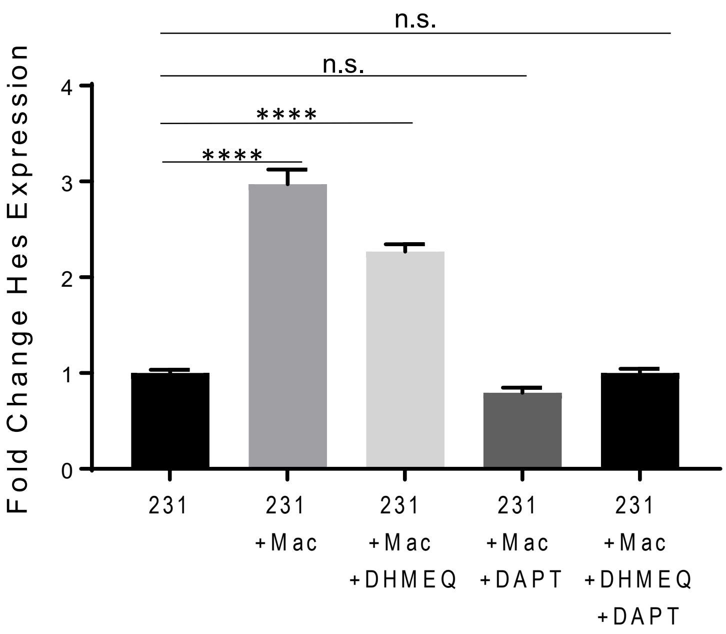

**Supplemental Figure 3.**

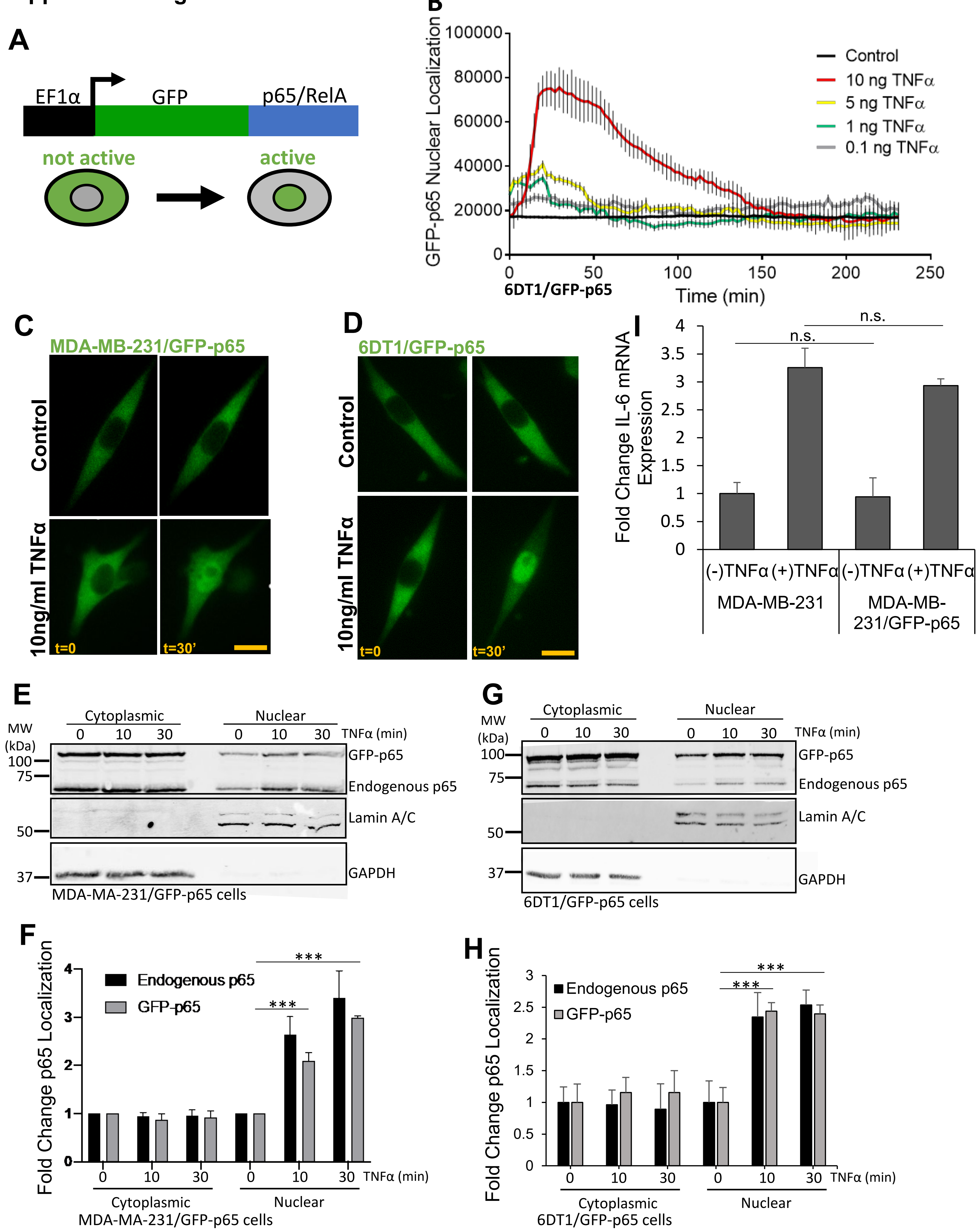

**Supplemental Figure 4.**

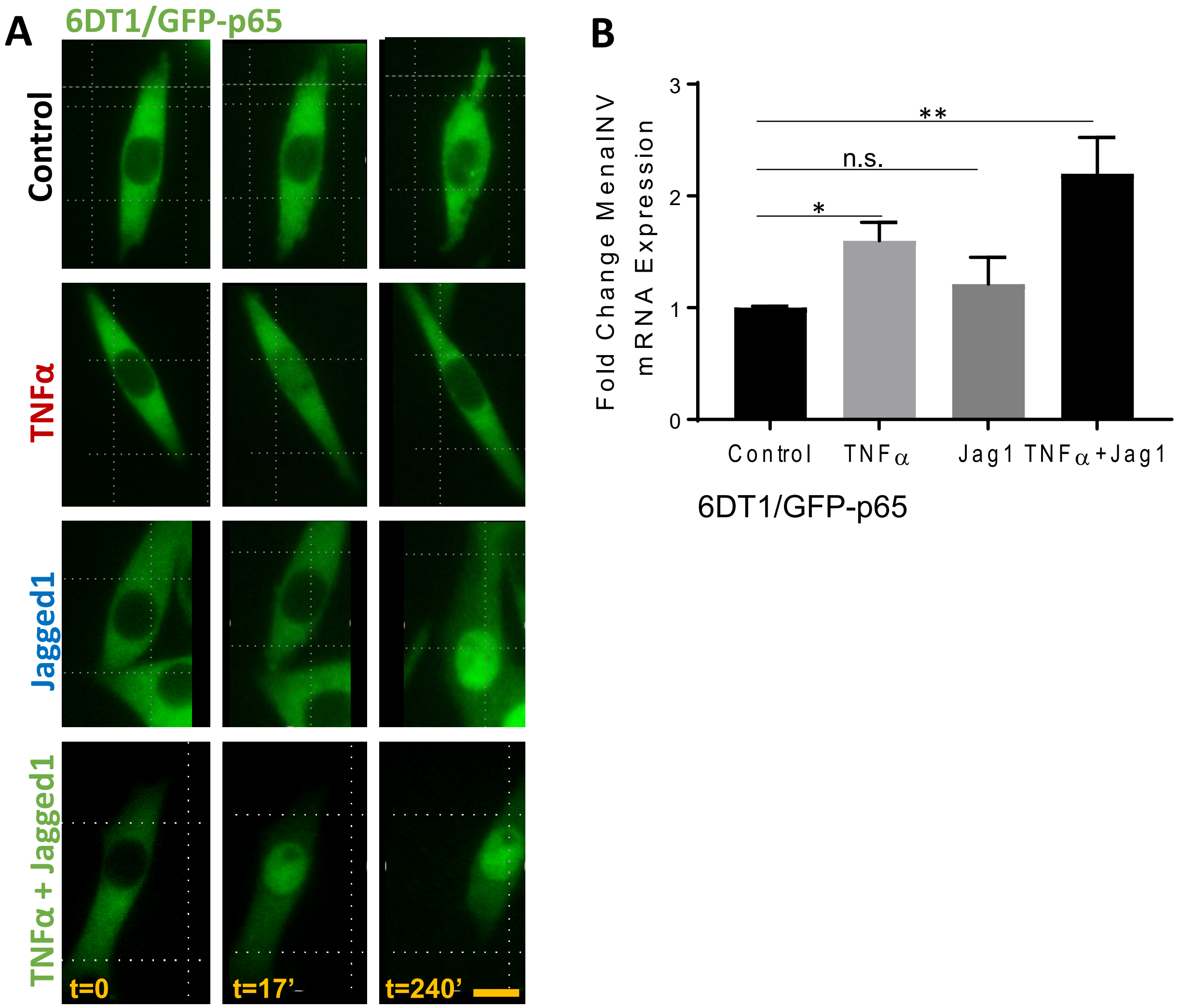

**Supplemental Figure 5**

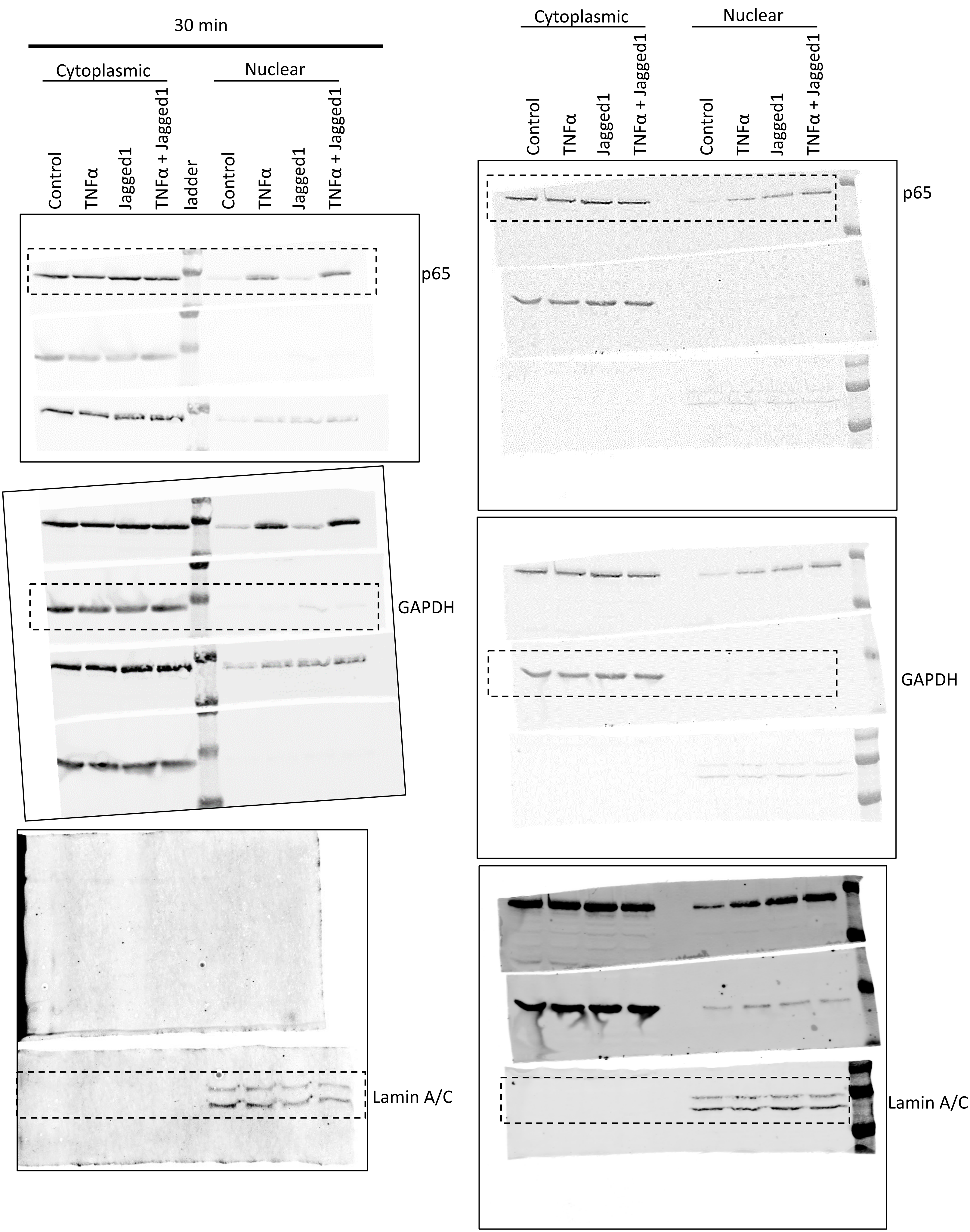

Supplemental Figure 6.

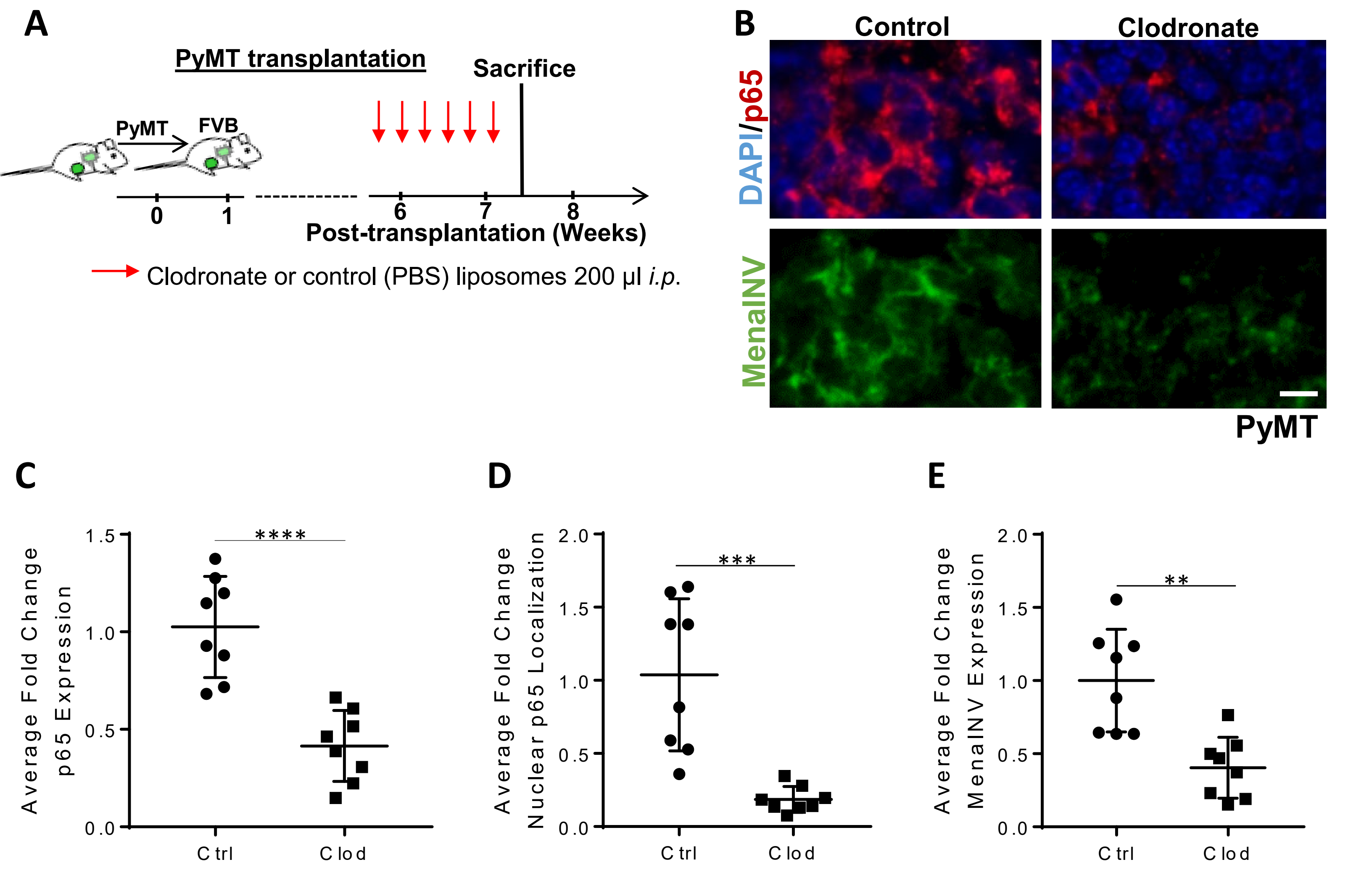

Supplemental Figure 7.

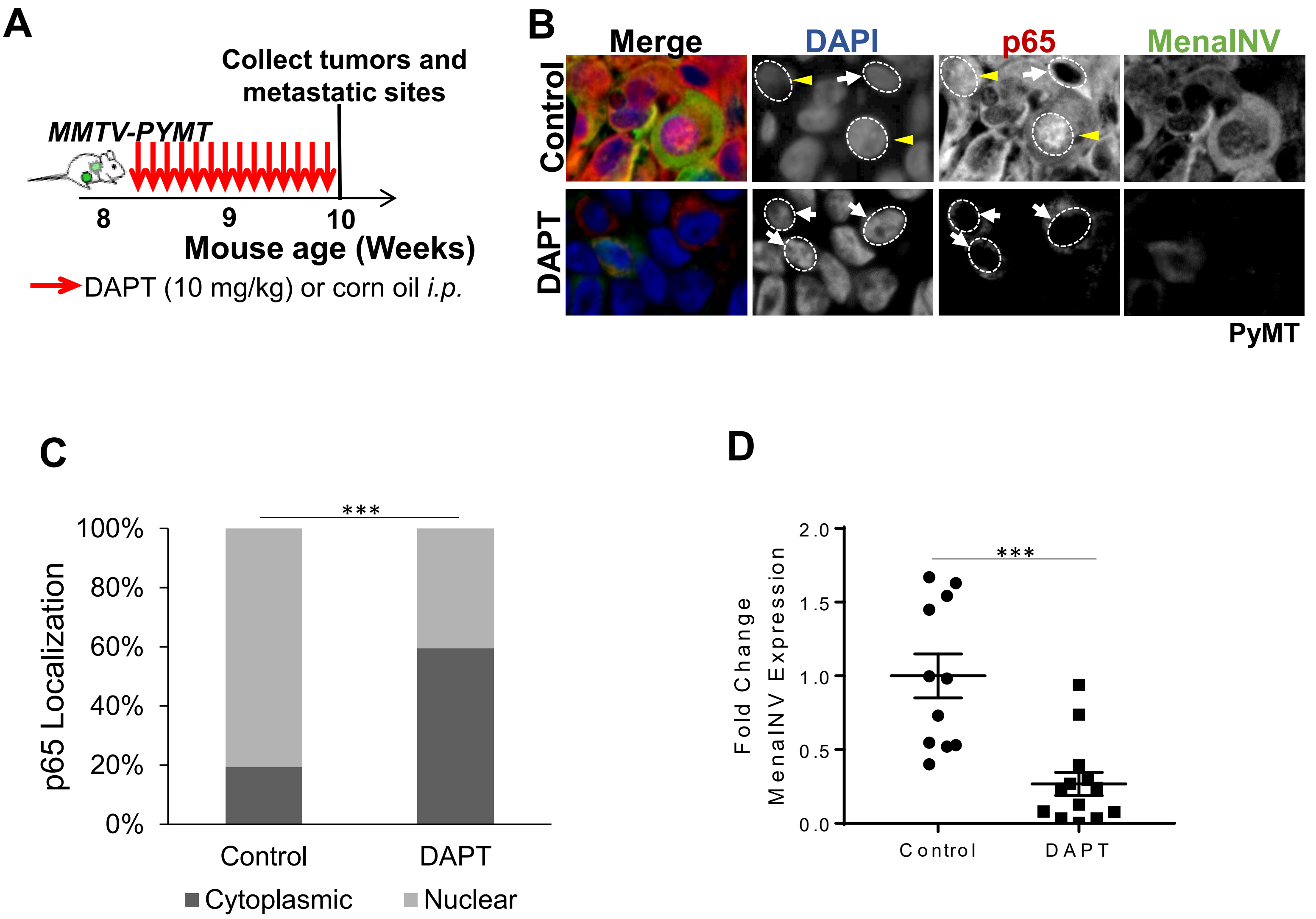

Supplemental Figure 8.

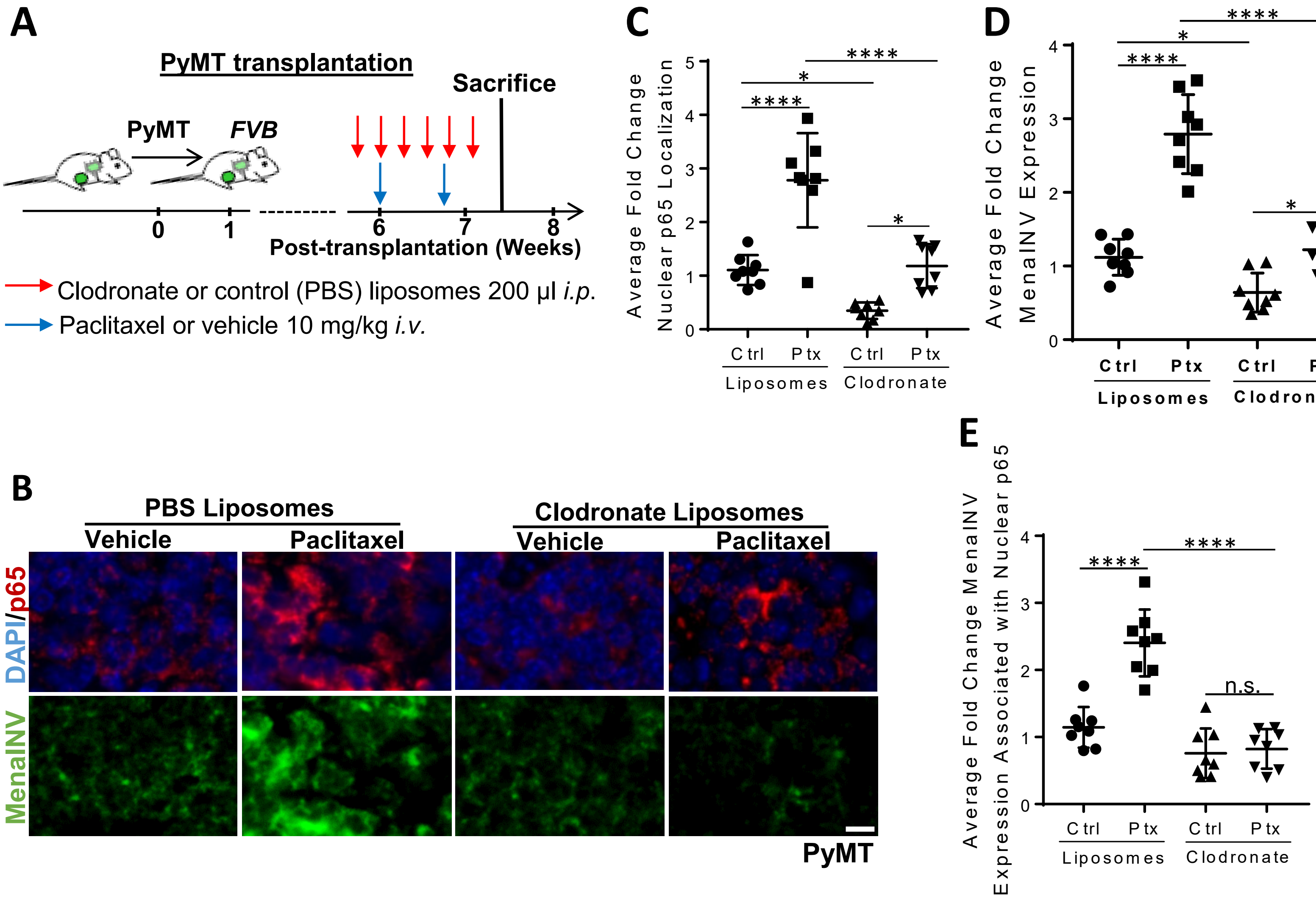

### Supplemental Figure Legends

**Supplemental Figure 1. The promoter sequences in the *ENAH* gene are capable of inducing Mena expression in breast cancer cells. (A)** The transcription factor binding sites found within 1,200 bp from the transcriptional start site in the promoter region of the *ENAH* gene across four species. The sequence of the κB binding site found in the human *ENAH* gene is noted. **(B)** MenaINV mRNA expression in MDA-MB-231 (231) cells treated with 0, 5, or 10 ng/ml active TGFβ for 24 hours. **(C)** MenaINV mRNA expression in MDA-MB-231 (231) cells treated with 0, 25 or 50 mM LiCl, which activates Wnt/B-catenin signaling, for 24 hours. The bars in (B) and (C) represent average fold change MenaINV mRNA compared to control (231 cells), +/- S.D. The data were analyzed using a one-way ANOVA with Dunnett's multiple comparisons test. n.s.=not significant.

**Supplemental Figure 2. Macrophage co-culture with MDA-MB-231 cells induces Hes1 expression.** mRNA expression of Hes (Notch1 transcriptional target) from extracts of MDA-MB-231 (231) cells cocultured with or without macrophages (Mac), the NF-κB inhibitor (DHMEQ), or Notch/gamma secretase inhibitor (DAPT) for 4 hours. The bars represent average fold change MenaINV mRNA compared to control (231 cells), +/- S.D. The data were analyzed using a one-way ANOVA with Tukey's multiple comparisons test. ****$p<0.0001$, ***$p<0.001$, n.s.=not significant.

**Supplemental Figure 3. NF-κB reporter (GFP-p65) is functional *in vitro*. (A)** Schematic of GFP-p65 reporter where GFP is linked to the N-terminus of full length human p65 (RelA), driven by an EF1α promoter. When the GFP-p65 reporter is localized primarily to the cytoplasm, the NF-κB signaling pathway is inactive. When the GFP-p65 reporter is nuclear (and can also be partially cytoplasmic), the NF-κB signaling pathway is active. **(B)** 6DT1/GFP-p65 tumor cells were treated with increasing doses of mouse TNFα (0 (control)- 10 ng/ml). Cells were imaged live for 240 minutes using an EPI fluorescence microscope for the duration of the treatment, with one captured image every 2.5 minutes. Quantification shows intensity of GFP-p65 nuclear localization over time following treatment. **(C, D)** The functionality of GFP-p65 reporter expressed in (C) MDA-MB-231 and (D) 6DT1 cells. The cells were treated with either vehicle or 10 ng/ml human TNFα for thirty minutes and time-lapse imaged live using an EPI fluorescence microscope for the duration of treatment. Stills from movies are shown at 0 and 30 minutes of treatment. Note the nuclear translocation of GFP-p65 following 30 minutes TNFα treatment. Scale bar = 10µm. **(E, G)** Nuclear and cytoplasmic fractions of (E) MDA-MB-231/GFP-p65 cells or (G) 6DT1/GFP-p65 cells treated with 10 ng/ml TNFα for 0, 10, or 30 minutes. Cell extracts were separated and western blotted using antibodies against p65, lamin A/C, and GAPDH. Note, the GFP-p65 reporter runs 30 kDa higher than the endogenous p65, at 65 kDa. **(F, H)** Quantification of western blots shown in (E and G) where the cytoplasmic and nuclear fractions from (F) MDA-MB-231/GFP-p65 or (H) 6DT1/GFP-p65 cells. p65 signal was normalized to the GAPDH and Lamin A/C loading controls, respectively. The 10- and 30-minute time points were set relative to the 0 minute time points to determine average fold change p65 expression, +/- SEM. **(I)** mRNA expression of IL-6 (NF-κB target gene) in MDA-MB-231 (control) or MDA-MB-231/GFP-p65 cells treated with vehicle or 10ng/ml TNFα for 4 hours. Bars show average fold change IL-6 mRNA expression of MDA-MB-231 cells treated with TNFα compared to untreated cells. Data in (D) and (F) were analyzed using a one-way ANOVA with Tukey's multiple comparisons test. ***p<0.001, n.s.=not significant.

**Supplemental Figure 4. Notch1 signaling enhances NF-κB signaling (sustained p65 nuclear localization).** Stills from movies at 0, 17, and 240 minutes of 6DT1/GFP-p65 cells treatment with vehicle, or 10 ng/ml human TNFα, or 80 µm Jagged1, or 10 ng/ml TNFα and 80 µm Jagged1. In all treatment groups with TNFα, the cells were treated for an initial 10 minutes, and then TNFα was washed out and replaced with minimal media, or with Jagged1 supplemented media. Cells were imaged live for 240 minutes using an EPI fluorescence microscope for the duration of the treatment, with one image captured every 2.5 minutes. Scale bar = 10 μm. **(B)** MenaINV mRNA expression in wild type MDA-MB-231 (231) cells treated as in (C) for 1 or 4 hours. Bars in (D) show average fold change MenaINV mRNA expression compared to Control at 1 or 4 hours. Data in (D) were analyzed using a one-way ANOVA with Tukey's multiple comparisons test. *$p<0.05$, **$p<0.01$, n.s.=not significant.

**Supplemental Figure 5. Full, uncropped western blotting images from Figure 2C showing Notch1 signaling enhances NF-κB signaling by sustaining p65 nuclear localization.** Full, uncropped western blot images showing the amount of p65 in the cytoplasmic and nuclear fractions of wild type MDA-MB-231 cells treated for 30 minutes (left blots) or 4 hours (right blots) with vehicle, or 10 ng/ml TNFα, or 80 μm Jagged1, or 10 ng/ml TNFα and 80 μm Jagged1 (TNFα + Jagged1). In all treatment groups with TNFα, the cells were treated for an initial 10 minutes, and then TNFα was washed out and replaced with minimal media, or with Jagged1 supplemented media. Dashed boxes display how the images where cropped for the main figure (Fig 2C).

**Supplemental Figure 6. Macrophage depletion decreases NF-κB signaling and MenaINV expression in PyMT transplantation model in vivo. (A)** Experimental design for macrophage depletion in PyMT transplantation model in FVB mice. *i.p.* = intraperitoneal. Red arrows indicate treatment days. **(B)** Immunofluorescence co-staining of the PyMT primary tumors from mice treated as outlined in (A) for p65 (red), MenaINV (green) and nuclei (blue-DAPI). Scale bar = 100μm. **(C)** Quantification of average fold change in p65 expression in PyMT transplanted mice from (A). **(D)** Quantification of average fold change in p65 nuclear localization in PyMT transplanted mice from (A). Only p65 co-localized with the nuclear DAPI signal was quantified.

**(E)** Quantification of average fold change MenaINV expression in PyMT transplanted mice from (A). Data in (C-E) were analyzed using a student's *t*-test. **p<0.01, ***p<0.001.

**Supplemental Figure 7. Inhibition of Notch1 signaling *in vivo* decreases activation of NF-κB signaling in the PyMT transplantation model. (A)** Schematic of DAPT treatment of *MMTV-PyMT* mice bearing transplanted breast tumors. Mice were treated with 10 mg/kg DAPT or vehicle (corn oil) by *i.p.* every day for 14 days. Red arrows represent treatment days. **(B)** Immunofluorescence staining of primary tumor tissues sections for DAPI (nuclear stain, blue), p65 (red) and MenaINV (green). White dotted circles indicate nuclei in the DAPI and p65 channels. Yellow arrow heads denote nuclei with p65 positive stain (active NF-κB signaling), white arrowheads indicate nuclei without p65 positive staining (inactive NF-κB signaling). **(C)** Quantification of p65 localization (% cytoplasmic/nuclear) in tumor tissue from (B). **(D)** Quantification of average fold change in MenaINV expression compared to control mice from (B). Data in (C) and (D) were analyzed using a student's *t*-test. ***p<0.001.

**Supplemental Figure 8. Chemotherapy treatment enhances NF-κB activation and MenaINV expression through macrophage recruitment in PyMT transplantation model. (A)** Experimental design of chemotherapy and clodronate treatments in PyMT transplantation model in FVB mice. *i.p.* = intraperitoneal, *i.v.* = intravenous. **(B)** Immunofluorescence staining of primary breast tumor tissues from mice treated as outlined in (A) with DAPI (nuclear stain, blue), and antibodies recognizing p65 (red), and MenaINV (green). **(C)** Quantification of average fold change in p65 nuclear localization in treated primary tumors from (A) stained for p65 and DAPI. Only p65 which co-localized with the nuclear DAPI signal was quantified. **(D)** Quantification of average fold change in MenaINV expression in treated primary tumors from (A). **(E)** Quantification of average fold change MenaINV expression associated with nuclear p65 staining of primary tumors from (A) stained for MenaINV. Data in (C-E) were analyzed using a one-way

ANOVA with Tukey's multiple comparisons test. *$p<0.05$, ***$p<0.001$, ****$p<0.0001$, n.s.=not significant.

**Supplemental Movies 1-8. Notch1 signaling sustains NF-κB over time.** Videos captured from **(1-4)** MDA-MB-231/GFP-p65 and **(5-8)** 6DT1/GFP-p65, cells treated with either (movies 1, 5) control or (movies 2, 6) 10 ng/ml TNFα, (movies 3, 7) 80 μm Jagged1, or (movies 4, 8) 10 ng/ml TNFα and 80 μm Jagged1 (TNFα+Jagged1). Cells were imaged live for 240 minutes using an EPI fluorescence microscope for the duration of the treatment, with one captured image every 2.5 minutes.
